# Dynamic changes in mRNA isoform usage during human retinal organoid development

**DOI:** 10.1101/2025.11.26.689062

**Authors:** Casey J. Keuthan, Sowmya Parthiban, Yen-Yu Chang, Xiaoqian Shan, Xiaoli Chang, Ethan Yan, Sheridan Cavalier, Winston Timp, Stephanie C. Hicks, Donald J. Zack

## Abstract

**Background:** Alternative mRNA splicing is a key mechanism for generating isoform diversity in eukaryotic cells. However, the extent of the splicing changes that occur during complex regulatory processes like neurodevelopment are still incompletely characterized.

**Results:** We performed nanopore-based long-read RNA sequencing on differentiating human stem cell-derived retinal organoids to identify temporal patterns of isoform usage across developmental stages. We found that retinal organoids undergo dynamic shifts in isoform usage throughout differentiation, which were not necessarily accompanied with changes in overall gene expression, as was observed for many genes involved in the regulation of mRNA splicing itself. Further analysis of human stem cell-derived retinal ganglion cells uncovered neuron-specific splicing signatures. Additionally, allele-specific gene expression analysis revealed extensive allelic imbalance in induced pluripotent stem cell-derived organoid cultures.

**Conclusions:** By combining direct long-read RNA sequencing with human stem cell retinal models we were able to develop a comprehensive database of isoform-level changes in differentiating human retinal cells during development. These results uncovered dynamic shifts in transcript usage during retinal differentiation, adding to our knowledge base of post-transcriptional RNA processing in the developing central nervous system and human *in vitro* culture systems.

## Background

The retina is a highly heterogenous central nervous system tissue that, during retinogenesis, undergoes a series of overlapping, strictly controlled processes to generate its diversity of cell types. High-throughput transcriptomics, especially single cell RNA-sequencing, has greatly shaped our understanding of transcriptional regulation of mammalian retinal cell development (1–6). However, our knowledge of how other, post-transcriptional mechanisms contribute to retinal cell differentiation is far more limited.

Alternative mRNA splicing is a fundamental post-transcriptional mechanism of gene regulation that enhances protein diversity in cells. Alternative splicing is especially high in neuronal tissues (7–10), where it contributes to many important physiological functions within the cells, including tightly regulated and complex processes like neurogenesis and synaptogenesis (11). Likewise, prior studies have shown that alternative splicing, including many retinal-specific splicing events such as cassette exon inclusion, as well as the use of alternative promoters, is frequent in the neuroretina (12–14). Moreover, not only can pathogenic variants in specific genes disrupt proper splicing and result in retinal disease, but mutations in several genes encoding ubiquitously expressed pre-mRNA processing factors themselves can preferentially result in retina-specific degeneration (15–17).

The emergence of third generation long-read sequencing technologies, like those of Oxford Nanopore Technologies (ONT) and PacBio, have greatly expanded our ability to examine isoform-level transcriptomic changes through real-time sequencing of entire transcripts of virtually any length, providing a more comprehensive landscape of the transcriptome with better resolution of individual isoforms and alternative splicing events. Recently, long-read RNA-sequencing has been used to profile isoform expression of cell surface markers in the developing mouse and adult human retina (18), as well as in multi-omics studies using post-mortem human retinal tissue (19), and short- and long-read sequencing approaches have been integrated in single cell analysis of the mouse retina (20). In contrast to these studies, isoform-level studies of developing human retinal cells have remained limited. Retinal organoids derived from human stem cells recapitulate much of the spatiotemporal development of the human retina, allowing us to examine developmentally regulated mRNA splicing events in the context of a human system. Moreover, the increasing popularity of using retinal organoids for drug screening and modeling retinal disease (21–23) further supports a pressing need for a more complete understanding of isoform-level transcriptomics in this *in vitro* model.

Here, we have generated one of the first long-read sequencing datasets for the developing human retina using direct cDNA ONT sequencing of stem cell-derived retinal organoids spanning multiple stages of retinogenesis. We identified temporally regulated isoform switching events across developmental time points, including those occurring in known retinal disease genes and genes involved in the regulation of mRNA splicing. We further show that isoform switching often occurs without significant changes to a gene’s overall expression, revealing transcript-level changes that would otherwise be overlooked with traditional differential gene expression analyses. We also identified neuron-specific splicing events through long-read sequencing of purified stem cell-derived retinal ganglion cells (RGCs). Finally, we leveraged our long reads to examine allele-specific gene expression where we found striking differences in allele-specific expression (ASE) between pluripotent stem cell types. Together, these data contribute to a more comprehensive understanding of post-transcriptional regulation in human retinal development.

## Results

### Long-read sequencing of developing retinal organoids

Many protocols have been established for differentiation of human stem cells into three-dimensional (3D) retinal organoids (24–28). With these, retinal organoid differentiation can be generally divided into three stages (Stages 1-3) that closely follow the same temporal pattern as *in vivo* human retinogenesis (29). Stage 1 (∼Weeks 4-8) organoids have a laminated morphology consisting of retinal progenitor cells majorly in the clear, outermost layer, while the center of the organoid is comprised of the early-born RGCs. During Stage 2 (∼Weeks 8-21), a subset of progenitor cells become further committed to a particular cell fate, such as rod and cone photoreceptor precursors, and by Stage 3 (∼Week 21+) the organoids give rise to a greater diversity of terminally differentiated retinal neurons and Müller glia cells. Organoids spanning each of these differentiation stages were separately pooled by time point for direct cDNA ONT sequencing to identify temporally regulated alternative splicing changes that occur during retinogenesis (**Figure 1A, Table S1**). On average, 99.4% of the reads per sample had primary mapped alignments to the reference genome with a Q score ≥ 10. The number of aligned ONT reads varied between ∼13-28 million for the different organoid stages, with a grand mean of 20.1 million total reads across all libraries (**Figure 1B**). Notably, ONT sequencing yielded consistent median N50 read lengths, averaging 2.3 kilobases (kb) across all libraries (**Figure 1C, S1, Table S2**). Nearly half of the mapped reads (45.7 ± 5.1%) spanned at least two exon-exon junctions, while over a fourth of reads (26.1 ± 4.0%) spanned four or more consecutive junctions (**Figure 1D**).

**Figure 1.**
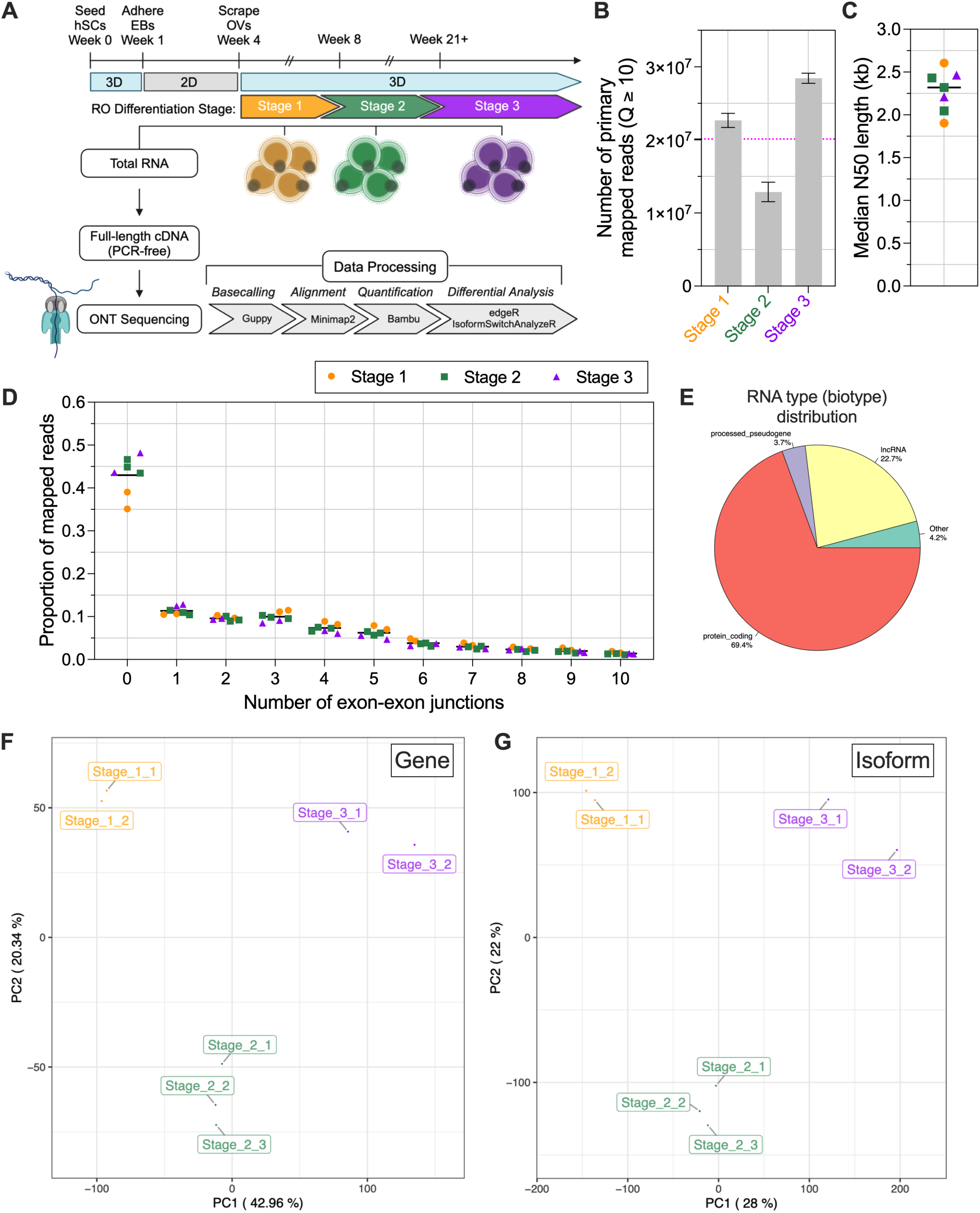
Direct cDNA ONT sequencing of human stem cell-derived retinal organoids. **A)** Experimental design and data analysis pipeline for long-read sequencing of human stem cell (hSC)-derived retinal organoids (ROs). Organoids were differentiated from hSCs aggregated into 3D embryoid bodies (EBs) over the course of the first week of cell culture, followed by a three-week adherent phase and subsequent scraping of the emerged optic vesicles (OVs) for continued differentiation as free-floating ROs. Total RNA was extracted from pooled sets of organoids at three stages of differentiation (Stages 1-3) for cDNA synthesis and direct ONT sequencing of the full-length cDNA libraries. Basecalling of ONT reads was performed with Guppy, followed by alignment with minimap2. Genes and isoforms were quantified using Bambu for subsequent differential expression and usage analysis, as well as visualization. Created with BioRender.com. **B)** Number of primary mapped ONT reads with a Q score ≥ 10 for each organoid stage. Error bars denote standard deviation. On average, 20.1 million primary mapped reads were obtained across the organoid samples (dashed pink line). **C)** Sequence length (as N50) for each organoid sample. The average N50 length was ∼2.3 kilobases (kb) amongst all organoid time points and replicates (black line). **D)** Histogram showing the proportion of primary mapped reads spanning different numbers of exon-exon junctions for each organoid sample. Line indicates grand mean across all organoid samples. **E)** RNA type (as biotype classification) of all detected transcripts. **F)** Gene and **G)** isoform expression level Principal Component Analysis (PCA) of the ROs.

Both known and novel transcripts were quantified from the aligned reads, which identified a total of 70,341 genes (including both protein- and non-protein-coding) and 278,023 isoforms (**Figure S2**). Most of the detected transcripts were classified as protein-coding (69.4%), though long non-coding RNAs (lncRNAs) and other types of transcripts were also found in the data (**Figure 1E**). We additionally filtered counts based on RNA type (according to ENSEMBL biotype), keeping protein-coding genes and filtering out known and novel isoforms in non-protein-coding genes. Furthermore, we filtered out genes and isoforms with low counts in edgeR (gene: *min.count* = 10, isoform: *min.count* = 2), which retained 16,971 genes and 55,708 isoforms for differential analyses (**Figure S2**). Principal Component Analysis (PCA) on both gene and isoform filtered counts clearly clustered the organoid samples based on differentiation stage (**Figures 1F-G**).

In addition to quantification of known transcripts, 225 novel genes and 1,118 novel isoforms were also identified in our pre-filtered data (**Figure S2**). Given that accurate prediction of novel transcripts remains a challenge (30), we also performed novel transcript discovery with IsoQuant to corroborate the novel transcripts detected by Bambu. We excluded all novel genes (IsoQuant detected 11,814 novel genes and 24,292 novel isoforms) and kept the 496 novel isoforms that were identified by both transcript discovery tools (**Figure S2)**. After additional transcript filtering, 342 novel isoforms mapping to 321 unique protein-coding genes were retained and included in our downstream analyses (**Table 1**).

### Long-read sequencing captures gene expression changes during retinal organoid differentiation

Short-read RNA sequencing has contributed significantly to our understanding of the gene level transcriptomic changes that occur during retinal development (1). To assess the reliability of long-read sequencing on stem cell-derived retinal organoids, we examined the gene-level concordance of our long-read data to comparable short-read methodology. Agarwal et al. recently generated a robust short-read (Illumina) bulk RNA sequencing dataset with time points ranging from as early as stem cell aggregation and spanning the three retinal organoid differentiation stages that underwent long-read sequencing in this study (for this comparison it should be noted that there can be significant differences between organoids generated by different labs, even when following similar protocols) (31). Overall, gene expression of the ONT sequencing samples correlated well with the short-read samples at comparable stages (**Figure 2A**). Correlation was high between Stage 1 ONT and both D25 and D65 short-read samples (Spearman correlation coefficients = 0.88 – 0.89), while Stage 2 ONT organoid samples correlated well with D65 and D100 short-read time points (Spearman correlation coefficients = 0.88 – 0.90). Stage 3 ONT samples were slightly less correlated with the short-read samples, but more closely matched later differentiation time points (≥ D65).

**Figure 2.**
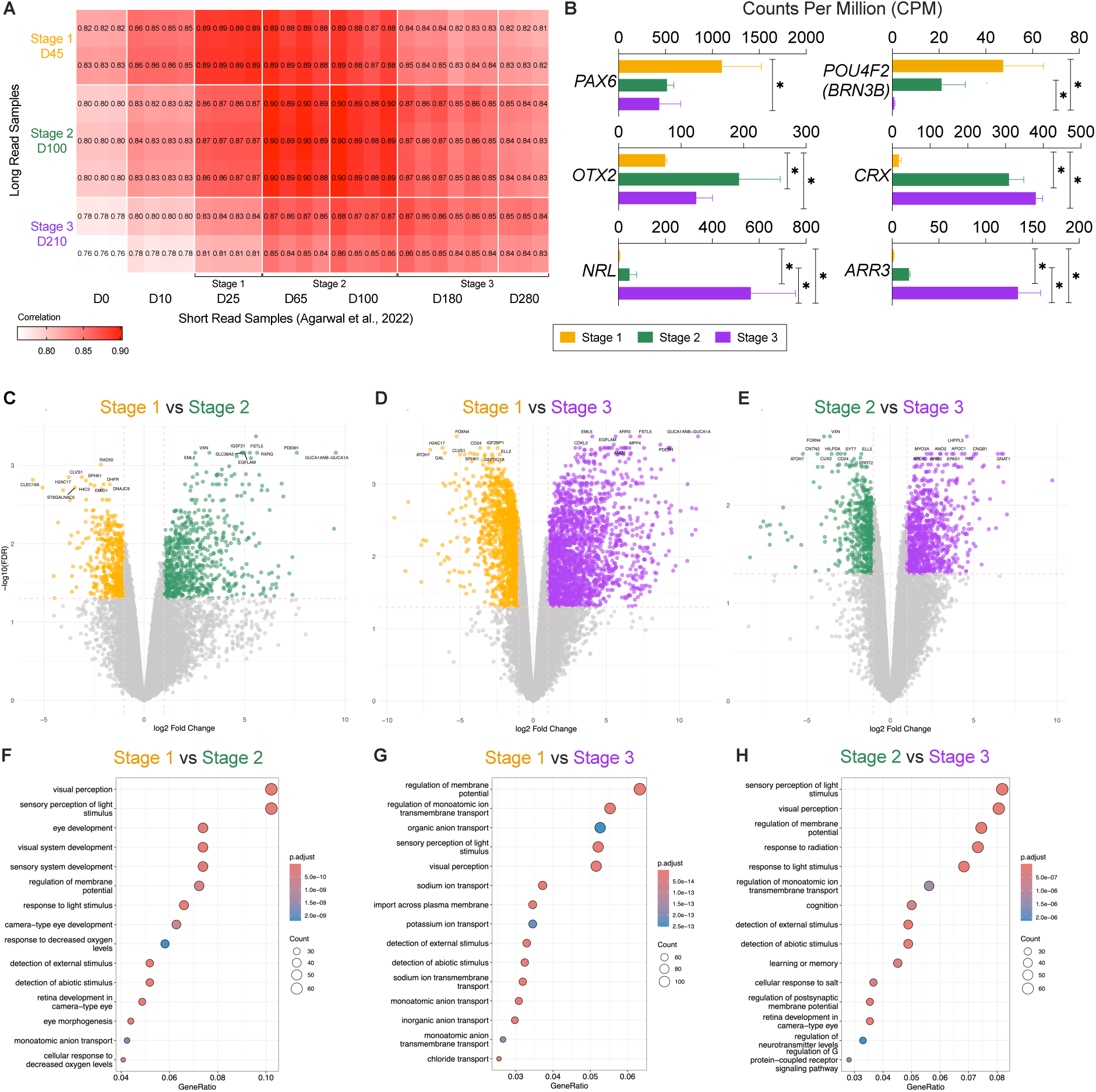
ONT sequencing captures gene-level transcriptomic changes in developing retinal organoids. **A)** Heatmap of Spearman’s rank correlation coefficients of raw gene Counts Per Million, (CPM) comparing retinal organoid differentiation time points from each direct cDNA ONT sequencing replicates and analogous short-read (Illumina) replicates from Agarwal et al. (31). **B)** Gene-level expression (plotted as CPM) from long-read sequencing replicates of common markers representing different cell types found at each retinal organoid differentiation stage. Stage 1: *PAX6* – paired box 6 (an eye field transcription factor, retinal progenitor cells); Stages 1-2: *POU4F2 (BRN3B)* – POU class 4 homeobox 2 (retinal ganglion cells); Stages 2-3: *OTX2* – orthodenticle homeobox 2 (photoreceptor and bipolar progenitor cells) and *CRX* – cone-rod homeobox (rod and cone photoreceptor precursor cells); Stage 3: *NRL* – neural retina leucine zipper (early rod photoreceptors) and *ARR3* – arrestin 3 (cone photoreceptors). Error bars indicate standard deviation. Asterisks denote gene is differentially expressed between groups (|log2FC| ≥ 1.0 & FDR < 0.05). **C-E)** Volcano plots highlighting differentially expressed genes between **C)** Stage 1 and Stage 2, **D)** Stage 1 and Stage 3, and **E)** Stage 2 and Stage 3 retinal organoids. Dashed lines indicate significance cut-offs used for differential gene expression analysis. **F-H)** Top gene ontology (GO) biological process terms from differentially expressed genes when comparing **F)** Stage 1 and Stage 2, **G)** Stage 1 and Stage 3, and **H)** Stage 2 and Stage 3 retinal organoid differentiation time points. Heatmap indicates adjusted p-value for enrichment, and dot size represents the number of significant genes found within each GO term.

We further investigated the extent to which biological (specific genetic pathways/programs) or technical (platform/methodology) differences may contribute to the genes driving the differences between the short- and long-read samples. DGE analysis between matched short- and long-read organoid samples did find more differentially expressed genes between Stage 3 ONT and D180 and D280 Illumina samples compared to matched Stage 1 and Stage 2 comparisons (**Table S3**). GO analysis of differentially expressed genes between the two platforms found an enrichment of cilia-related pathways in the long-read samples at earlier stages, while eye development/visual perception gene expression was higher for many genes in the Stage 3 long-read samples compared to the later short-read time points (**Figure S3**).

Conversely, at any stage, downregulated genes in the long-read samples (enriched in short-read) were largely dominated by metabolic pathways (**Figure S3**). Notably, there were a number of genes involved in axonogenesis that were less expressed in the Stage 2 long-read samples, possibly reflective of differences in the surviving populations of RGCs (which are known to die over time in 3D culture (26, 32)) between the two protocols and/or time points compared (**Figure S3**). At every differentiation stage, upregulated genes in the long-read samples had lower median GC content, consistent with the known GC-content bias in short-read/PCR-amplified sequencing methodologies (33, 34) (**Figure S4)**. While upregulated genes in the long-read samples also had longer transcript lengths compared to genes enriched in the short-read samples (downregulated in long-read) at all stages, this difference was most pronounced in the Stage 3 comparison (**Figure S4, Table S3**), which could be a contributing source to lower correlation with the short-read samples.

Within our long-read data, we found significant differences in gene expression between the different organoid stages (**Figure 2C-E**). Differential gene expression (DGE) was the greatest when comparing the earliest (Stage 1) and latest (Stage 3) differentiation stages, with a total of 4,144 genes differentially expressed (2573 upregulated, 1571 downregulated), while we found 1,220 and 1,633 differentially expressed genes between early (Stage 1 vs Stage 2) and later (Stage 2 vs Stage 3) developmental stages, respectively (Stage 1 vs 2: 730 upregulated, 490 downregulated; Stage 2 vs 3: 994 upregulated, 639 downregulated) (**Table S4**). Moreover, we could identify differential gene expression changes in known marker genes at each organoid differentiation stage in our long-read sequencing data (**Figure 2B, S5**). For example, early eye development genes like *PAX6* and *VSX2*, as well as early born cell-type markers such as RGC-specific *POU4F2* (*BRN3B*) and *SNCG,* were highly expressed in Stage 1 organoids, with markers associated with early neuronal/photoreceptor differentiation like *OTX2*, *CRX*, and *NEUROD1* increasing in Stage 2 organoids, and by Stage 3 there was significant expression of photoreceptor genes such as *NRL*, *ARR3*, *NR2E3*, *RCVRN*, and *RHO*. Finally, differentially expressed genes showed enrichment of expected biological processes related to retinogenesis, including “sensory perception of light stimulus” (GO: 0050953) and “visual perception” (GO: 0007601) (**Figure 2F-H**).

### Long-read sequencing detects greater isoform diversity in retinal organoids compared to short-read sequencing

The isoforms expressed amongst all organoid samples were associated with 15,683 unique genes (**Figure S2**). Over 69.9% of these genes encoded for two or more isoforms (**Figure 3A**). To more directly assess how this compared to traditional short-read sequencing, we also performed identical isoform quantification, this time using Salmon (35), on both our long-read samples and the short-read retinal organoid dataset from our earlier, gene-level analysis. Given the length constraints of Illumina reads, the short reads spanned considerably fewer exon-exon junctions than what we captured with direct ONT sequencing, as expected (**Figure 1D, S6**). Notably, under identical quantification, we could detect slightly more unique isoforms (≥ 1 count in ≥ 1 sample) in the long-read samples (226,256 isoforms across 45,533 genes in long-read and 203,124 isoforms across 58,248 genes in short-read). However, after filtering for protein-coding genes and minimum counts (edgeR, *min.count* = 2), we retained 103,949 isoforms across 17,585 genes in the long-read samples and 117,762 isoforms across 17,738 genes, which may be attributed to having four times as many samples and including a wider array of developmental time points (in which earlier, gene-level analysis showed lower correlation of our long-read data to the early short-read time points) in the short-read samples compared to our long-read dataset. Yet, when we compared the isoform distribution across genes for the two platforms, we found that we could detect a greater diversity of isoforms per gene in the long-read samples (detecting ≥ four isoforms per gene) (**Figure S6**). This was most apparent in our ability to detect genes with large numbers of known isoforms, where we found 34.8% more genes with over ten isoforms detected, on average (mean = 2,329 genes for long-read and 1,518 genes for short-read) (**Figure S6**). Finally, we examined the pathways that the 7,500 genes (from 15,316 isoforms) unique to our long-read dataset were associated with and found an enrichment of GTPase signal transduction and axonal pathways, which have relevance to retinal development and function (**Figure S6**).

**Figure 3.**
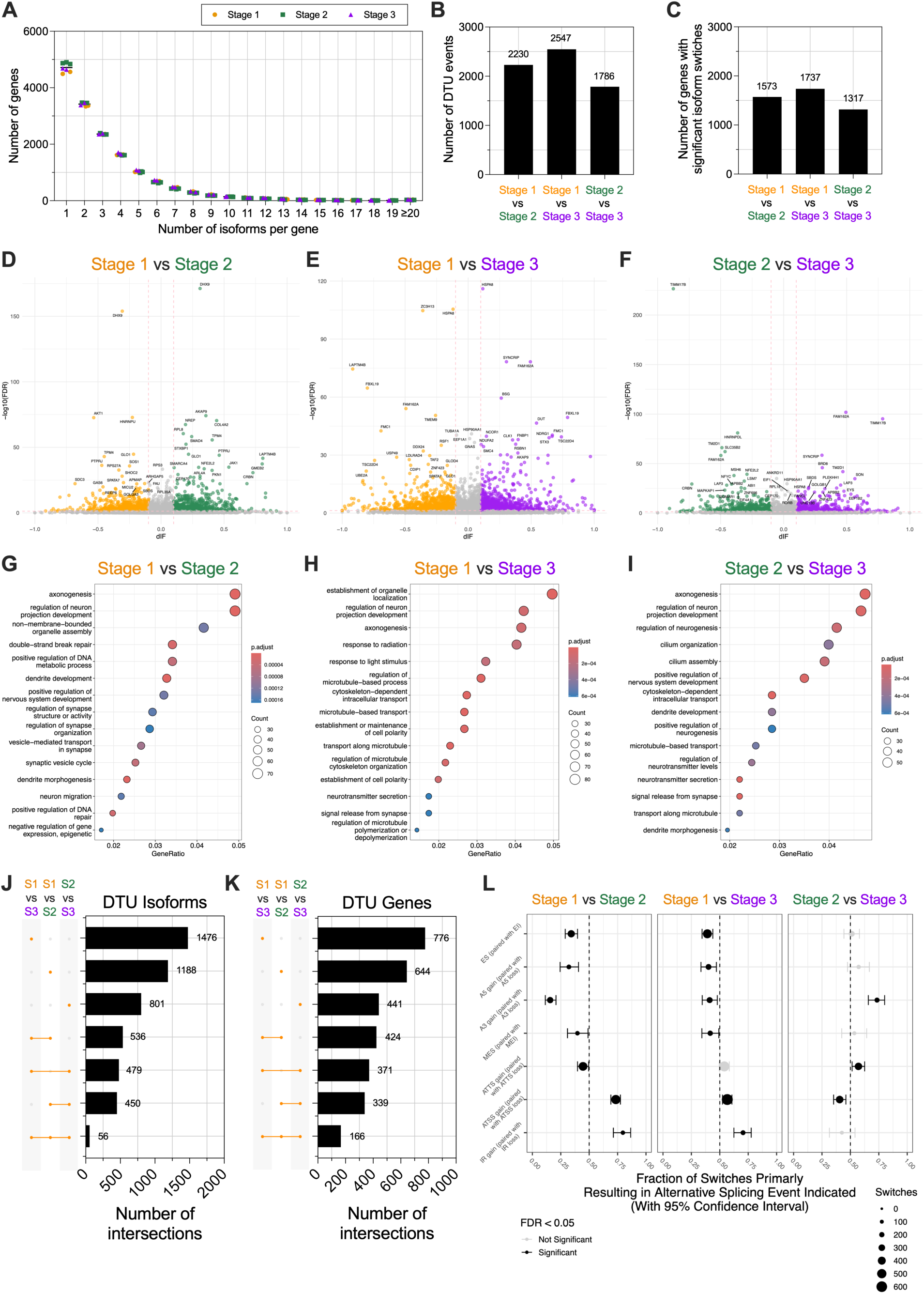
Isoform-level transcriptomic changes with retinal organoid differentiation. **A)** Histogram showing the number of isoforms detected per gene for each organoid sample. Line indicates overall mean across all organoid samples. **B)** Number of significant isoform switches (|dIF| ≥ 0.1 & qval < 0.05) occurring at different stages of organoid differentiation. **C)** Number of genes with significant isoform switches. **D-F)** Volcano plots highlighting differentially used transcripts between **D)** Stage 1 and Stage 2, **E)** Stage 1 and Stage 3, and **F)** Stage 2 and Stage 3 retinal organoids. Dashed lines indicate significance cut-offs used for DTU analysis. **G-I)** GO pathway analysis showing enriched biological processes for genes with significant isoform switches. Heatmap indicates adjusted p-value for enrichment, and dot size represents the number of significant genes found within each GO term. **J-K)** Upset plots showing the number of shared **J)** genes with significant isoform switches and **K)** differentially used isoforms between comparisons. **L)** Types and enrichment of alternative splicing events found in genes with significant isoform switches. Dot size indicates the number of significant switches. Red = significant skew of an alternative splicing event (qval < 0.05). ES/EI – exon skipping/inclusion; A5 – alternative 5’ splice site; A3 – alternative 3’ splice site; MES/MEI – multiple exon skipping/inclusion; ATTS – alternative transcription termination site; ATSS – alternative transcription start site.

### Developmentally controlled isoform switching in retinal organoids

We performed differential transcript usage (DTU) analysis across organoid stages to determine how many of genes undergo “isoform switching” during retinal organoid differentiation. DTU examines changes in the abundance of a gene’s different isoforms as a proportion of the overall expression of the gene (36, 37). Significant isoform switching was defined as a transcript having at least a 10% difference in its relative abundance between conditions (differential isoform fraction, |dIF| ≥ 0.1), along with an adjusted p-value (q-value) cut-off < 0.05. Our DTU analysis identified thousands of significant isoform switching events across differentiation stages (**Figure 3B**). We found 2,230 DTU isoforms between Stage 1 and Stage 2 organoids, with these isoform switches occurring in 1,573 unique genes (**Figure 3B-D**). There were slightly less DTU events when comparing Stage 2 organoids with later Stage 3 samples (1,786 DTU isoforms in 1,317 genes) (**Figure 3B-C, F**). Not surprisingly, the highest number of DTU events were found when comparing isoform usage between our earliest (Stage 1) and latest (Stage 3) organoid stages (2,547 DTU isoforms in 1,737 genes) (**Figure 3B-C, E**). Across all three comparisons, the average magnitude of difference in transcript usage for a significant event was ∼25% (mean |dIF|: Stage 1 vs 2 = 0.242, Stage 1 vs 3 = 0.258, and Stage 2 vs 3 = 0.249), while a small subset of DTU events had dIF values greater than 0.4 (12.8% of DTU events across all comparisons) (**Figure S7, Table S5**).

GO analyses on the genes that underwent isoform switching were less specific to the sensory/visual system as seen in the earlier differential gene expression analysis (**Figure 2F-H**) and instead included broader, pan-neuronal processes (**Figure 3G-I**). Genes with significant DTU in Stage 1 vs Stage 2 organoids were enriched in axon/synapse development-related processes, followed in later organoid stages by pathways associated with more mature neuronal characteristics like cell polarity, cilia development, and neurotransmission (**Figure 3G-I**). We also identified subsets of common DTU events between the different comparisons (**Figure 3J-K**). While many DTU isoforms and genes with isoform switches were unique to a particular stage/comparison, 166 genes had significant isoform switches regardless of the comparison. These isoform switching events predominantly occurred in genes related to the release of neurotransmitters. (**Figure S9**).

### Alternative splicing events are specific to developmental stage

We further investigated the types of alternative splicing events found in the developmentally controlled isoform switches. Cassette exon skipping/inclusion (ES/EI), as well as alternative transcription start and termination site (ATSS and ATTS, respectively) splicing, were the most abundant types of splicing events in DTU isoforms across all three comparisons (**Figure 3L**). Notably, there was uneven usage for many types of splicing events. This was most extensively found for Stage 1 vs Stage 2 switches where alternative splicing events were significantly skewed (e.g., more exon skipping than inclusion events), while the only significant bias in splice type usage in Stage 2 vs Stage 3 switches were towards loss of alternative 3’ splice sites (A3) and ATTS and gains in ATSS (**Figure 3L**). Interestingly, bias in splicing type usage differed when comparing earlier (Stage 1 vs Stage 2 or Stage 1 vs Stage 3) and later (Stage 2 vs Stage 3) differentiation stages. For example, A3 splice sites were significantly gained in isoform switches comparing Stage 1 organoids but shifted to A3 splice site loss in switches found between older organoids (**Figure 3L**).

These alternative splicing changes can confer various downstream consequences that affect transcript diversity in a cell/tissue. To understand the potential functional impacts of these isoform switches, we analyzed the predicted consequences of the significant DTU events across all organoid stage comparisons (**Figure S10**). We found that these switches significantly affected ORFs, as well as changes to protein domains, N-terminus signal peptides, and differential sensitivity to NMD (**Figure S10**). These functional consequences closely aligned with the alternative splicing changes found in our earlier analysis (e.g., longer/shorter or gain/loss of ORFs coupled with cassette exon splicing and intron retention), with significant differences in the enrichment and depletion pattern of these consequences in younger versus older retinal organoids (**Figure 3L, S10**).

### Isoform switching in retinal disease genes during organoid development

Retinal disease encompasses a heterogeneous collection of disorders characterized by progressive vision loss and blindness due to dysfunction and degeneration of the retina. To date, over 300 different genes and genetic loci have been associated with retinal disease (38). Retinal organoids have become an increasingly popular tool to model retinal disorders, whether for studying disease pathologies or screening drugs and other potential therapies in the context of human retinal cells. Thus, it is important to know the isoform usage pattern of retinal disease genes in this model system. We investigated how many genes associated with retinal disease (genes acquired from RetNet, the Retinal Information Network (38)) underwent significant isoform switching during organoid development (**Table 2**). As expected, gene-level expression of many retinal disease genes was significantly higher in Stage 3 retinal organoids (**Figure S11**); however, we found 82 retinal disease genes underwent significant isoform switching during organoid differentiation (**Figure 4A**). Notably, multi-isoform genes were overrepresented within the subset of retinal disease genes compared to all genes detected in our transcriptome-wide analysis (one-sided Chi-square test; odds ratio = 1.76, p-value = 0.0001) (**Figure S12, Table S6**). Among multi-isoform genes with significant DTU, retinal disease genes were also significantly enriched (one-sided Chi-square test; odds ratio = 1.59, p-value = 0.0005) (**Figure 4B, Table S6**). This is consistent with reports that the expansive isoform diversity within nervous system tissues regulates intrinsic neuronal properties (10), since many of these retinal disease-associated genes are preferentially expressed in the retina and are essential for proper development and function (39, 40).

**Figure 4.**
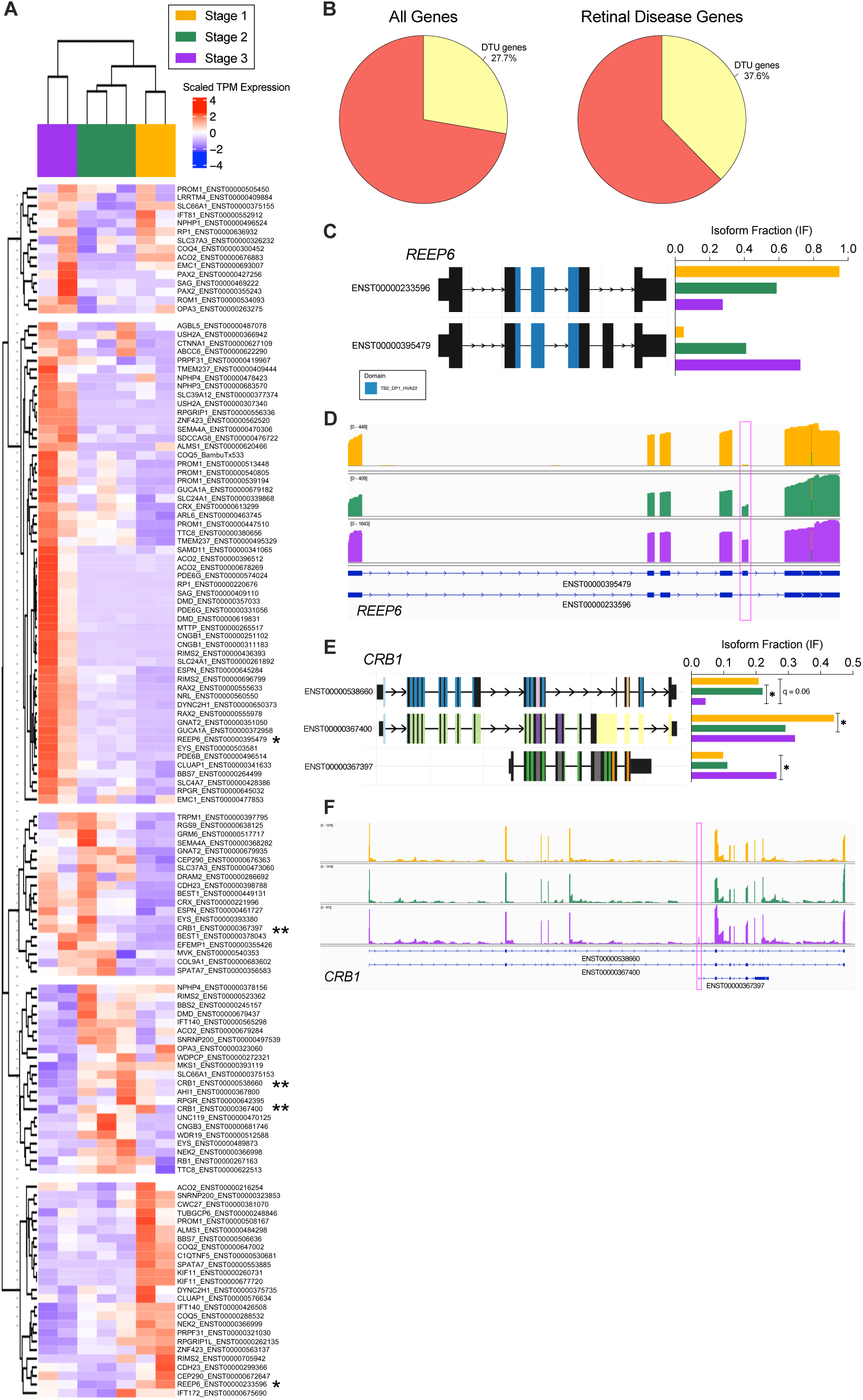
Temporally regulated isoform switching of retinal disease-associated genes. **A)** Heatmap of significant isoform switches in retinal disease genes. Colors indicate scaled expression (transcripts per million, TPM) of each isoform. Asterisks highlight DTU isoforms for *REEP6* (*) and *CRB1* (**). **B)** Proportion of genes with significant DTU in all genes (left) and within the subset of retinal disease genes (right). Percentages are calculated from the numbers of multi-isoform genes. **C)** *REEP6* (receptor accessory protein 6) protein-coding isoforms expressed (left) and isoform usage (as isoform fraction, IF) during different retinal organoid stages (right). Color exons denote conserved protein domain (TB2_DP1_HVAA22). **D)** IGV browser showing ONT reads mapping to specific *REEP6* isoforms. Pink box highlights alternatively spliced cassette exon. **E)** *CRB1* (crumbs cell polarity complex component 1) isoforms (left) and corresponding isoform fractions (right) for transcripts with significant DTU in retinal organoids. **F)** IGV browser showing reads mapping to different *CRB1* isoforms. Pink box highlights alternatively transcription start site for the ENST00000367397 variant.

One of these most striking isoform switches in our long-read dataset was for receptor expression enhancing protein 6 (*REEP6*), which has two protein-coding isoforms that differ through alternative splicing of a single cassette exon (**Figure 4C**). The retina, and more specifically rod photoreceptors, are known to preferentially express the *REEP6* variant in which this exon is included, and it has been demonstrated that *REEP6* alternative splicing is necessary for normal retinal function (41, 42). We found a clear decrease in usage of the exon-skipping variant in tandem with increased use of the photoreceptor-specific isoform over time, which correlates with the birth of rod photoreceptors in the organoids (**Figure 4C-D, S5**). We also found multiple crumbs cell polarity complex component 1 (*CRB1*) isoforms with usage patterns that varied across organoid stages (**Figure 4E-F**). These included two alternatively spliced, longer *CRB1* transcripts (ENST00000538660 and ENST00000367400), along with a shorter variant (ENST00000367397) predominantly used in Stage 3 retinal organoids (**Figure 4E-F**).

### Significant isoform switching in non-differentially expressed genes

As mentioned earlier, we detected significant gene-level expression changes through retinal organoid development and found that both differential gene and isoform expression changes were by far the greatest when comparing the earliest (Stage 1) and latest (Stage 3) organoid differentiation stages (**Figure 2D, 5A**). But it is important to consider that changes in usage between transcripts could be compensatory and may not result in changes in the overall expression of a gene, differences that would go otherwise undetected by conventional differential expression analyses. We found that isoform switching was largely unrelated to significant expression changes, particularly at the gene level (**Figure 5A-B**). This was most distinct in the Stage 1 vs Stage 2 organoids, where only 119 of the 1,573 genes with significant DTU were differentially expressed at this stage (**Figure 5B**). Even between Stages 1 and 3, where we observed the highest numbers of both DGE and DTU, we found less than 25% of the DTU genes had significant gene-level expression differences (**Figure 5B**). One of many examples includes basigin (*BSG*), which encodes a transmembrane glycoprotein with a known photoreceptor-specific splice variant (43, 44) (**Figure 5C-E**). *BSG* was expressed at high levels throughout organoid differentiation but did not exhibit differential gene expression (**Figure 5C**). However, we found significant differences in isoform usage between alternatively spliced cassette exon transcripts (**Figure 5D-E**).

**Figure 5.**
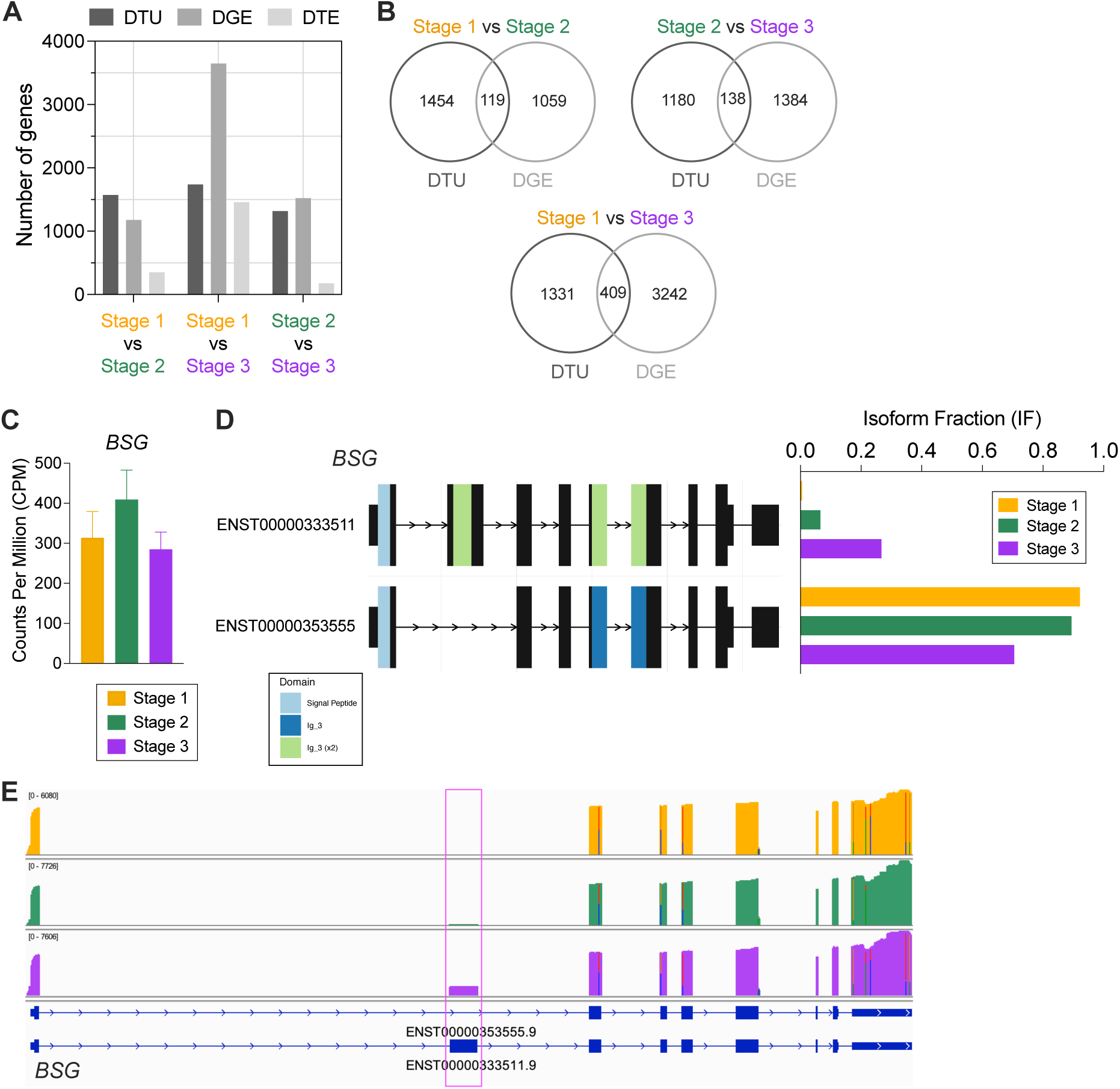
Dynamic isoform switching during organoid differentiation without gene expression changes. **A)** Number of genes with significant DTU (|dIF| ≥ 0.1 & qval < 0.05), DGE (|log2FC| ≥ 1.0 & qval < 0.05), and DTE (|log2FC| ≥ 1.0 & qval < 0.05). **B)** Overlapping genes differentially expressed and undergo significant isoform switching. **C-D)** *BSG* (basigin) **C)** gene expression plotted as CPM and **D)** visualization of the two expressed isoforms (left) and associated isoform usage (right) in differentiating retinal organoids (plotted as isoform fraction, IF). Colored exon regions denote conserved protein domains. Signal Peptide – Signal peptide domain, Ig_3 – Immunoglobulin domain 3. **E)** IGV browser showing reads mapping to specific *BSG* isoforms. Pink box highlights alternatively spliced cassette exon.

### Isoform-level regulation of splicing genes during retinal organoid development

We next sought to investigate the regulation behind the observed isoform switching. RNA splicing is a highly orchestrated process involving hundreds of splicing factors and RNA binding proteins (RBPs) in its regulation. Using the GeneCards database (45) we curated a list of over 500 genes involved in mRNA splicing and expressed in our long-read data to determine how expression of splicing-related genes correlated with the significant DTU events. This gene list included RBPs and spliceosome components (many of which are known to be ubiquitously expressed across all tissue and cell types), as well as known retinal-specific splicing factors (e.g., *MSI1/2*, *RP9*, *SRRM3*) (46–49) (**Table 3**). Many of these splicing-related genes were highly expressed and expressed throughout retinal organoid differentiation, though a small subset (102 genes) were differentially expressed between organoid stages (**Figure 6A-C**). These included genes like polypyrimidine tract binding protein 1 (*PTBP1*), a well-known negative regulator of splicing in neuronal differentiation (50–53), which decreased in later organoid stages (**Figure 6D**). Other retinal splicing factors, like *MSI2* and *SRRM3*, were also differentially expressed at different retinal organoid time points (**Figure 6D**).

**Figure 6.**
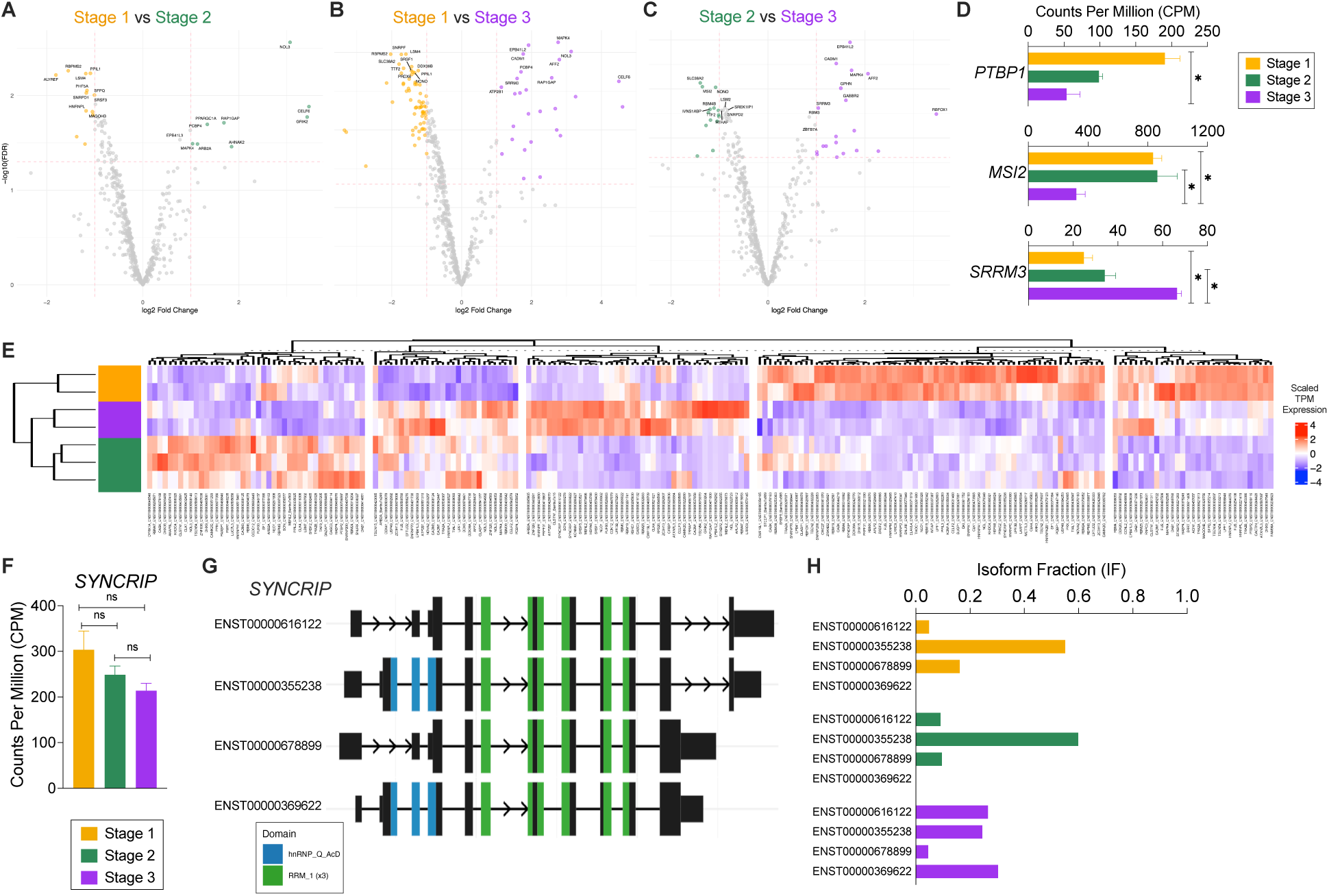
Splicing regulation during retinal organoid differentiation. **A-C)** Volcano plots highlighting differentially used transcripts for splicing-related isoforms comparing **A)** Stage 1 and Stage 2, **B)** Stage 1 and Stage 3, and **C)** Stage 2 and Stage 3 retinal organoids. Dashed lines indicate significance cut-offs used for DTU analysis (|dIF| ≥ 0.1 & qval < 0.05). **D)** Gene expression levels (as CPM) of the RNA-binding protein *PTBP1* (polypyrimidine tract-binding protein 1) and retina-enriched splicing factors *MSI2* (musashi RNA binding protein 2) and *SRRM3* (serine/arginine repetitive matrix 3). Error bars indicate standard deviation. Asterisks denote gene is differentially expressed between groups (|log2FC| ≥ 1.0 & qval < 0.05). **E)** Heatmap of isoform switching in splicing genes. **F)** *SYNCRIP* (synaptotagmin-binding cytoplasmic RNA-interacting protein) gene expression (plotted as CPM). Error bars represent standard deviation. ns = not significant. **G)** Visualization of significantly used *SYNCRIP* isoforms. Colors denote location of conserved protein domains encoded within the mRNA transcript; hnRNP_Q_AcD – Heterogeneous nuclear ribonucleoprotein Q acidic domain; RRM_1 – RNA Recognition Motif 1. **H)** Isoform usage of *SYNCRIP* transcripts plotted as isoform fraction (IF).

It is important to note that many of these splicing-related genes can also be alternatively spliced to generate multiple protein-coding mRNA isoforms. Despite few gene-level expression differences, we found many splicing-related genes also underwent isoform switching with organoid differentiation (**Figure 6E**). One example of this is synaptotagmin binding cytoplasmic RNA interacting protein (*SYNCRIP1*), an hnRNP family member associated with a number of neurological diseases, 456 including autism spectrum disorder and epilepsy (54). Whereas SYNCRIP gene expression was high but not significantly different across organoid time points (Figure 6F), SYNCRIP isoform usage patterns varied between younger and older samples (**Figure 6G-H**). In Stage 1 and 2 organoids, there was preferred usage of the ENST00000355238 isoform and no detection of ENST00000369622, while Stage 3 organoids showed equal usage of three SYNCRIP variants (**Figure 6G-H**). Together, these data further add to the complexity inherent in the regulation of mRNA splicing in neuronal development.

### Understanding mRNA splicing differences specific to retinal neurons

One limitation to bulk transcriptomics of retinal organoids is that, at any given stage of differentiation, there are vastly different populations and proportions of retinal progenitor cells and terminally differentiated neurons. RGCs are an early-born cell type found in Stage 1 retinal organoids; however, these RGCs eventually die in 3D culture (26, 32), which is reflected by the progressive reduction in RGC marker gene expression in the later stage organoids samples (**Figure 2B, S5**). To investigate isoform usage events in specific retinal neurons, we independently differentiated cells into RGCs using an established differentiation protocol (55), coupled with a CRISPR-based method to purify differentiated RGCs from the other cells in the 2D culture (**Figure 7A**) for long-read sequencing **(Figure S14, Table S8**). Our long-read data could detect a significant enrichment of RGC gene markers in the purified samples compared to unbound flow-through (FT) (**Figure 7B**). Transcript usage analysis between these groups identified 1,624 unique isoforms in 1,081 genes that had significant DTU (**Figure 7C-D**). These DTU genes were largely involved in pathways related to axonal transport and membrane-cytoskeletal interactions (**Figure 7E**). For example, the actin *ADD1* (adducin-1), a gene involved in the formation and stabilization of the actin cytoskeleton, is highly expressed in both RGCs and FT samples (**Figure 7F**) but had a significant difference in usage between two of its protein-coding isoforms (**Figure 7G-H**).

**Figure 7.**
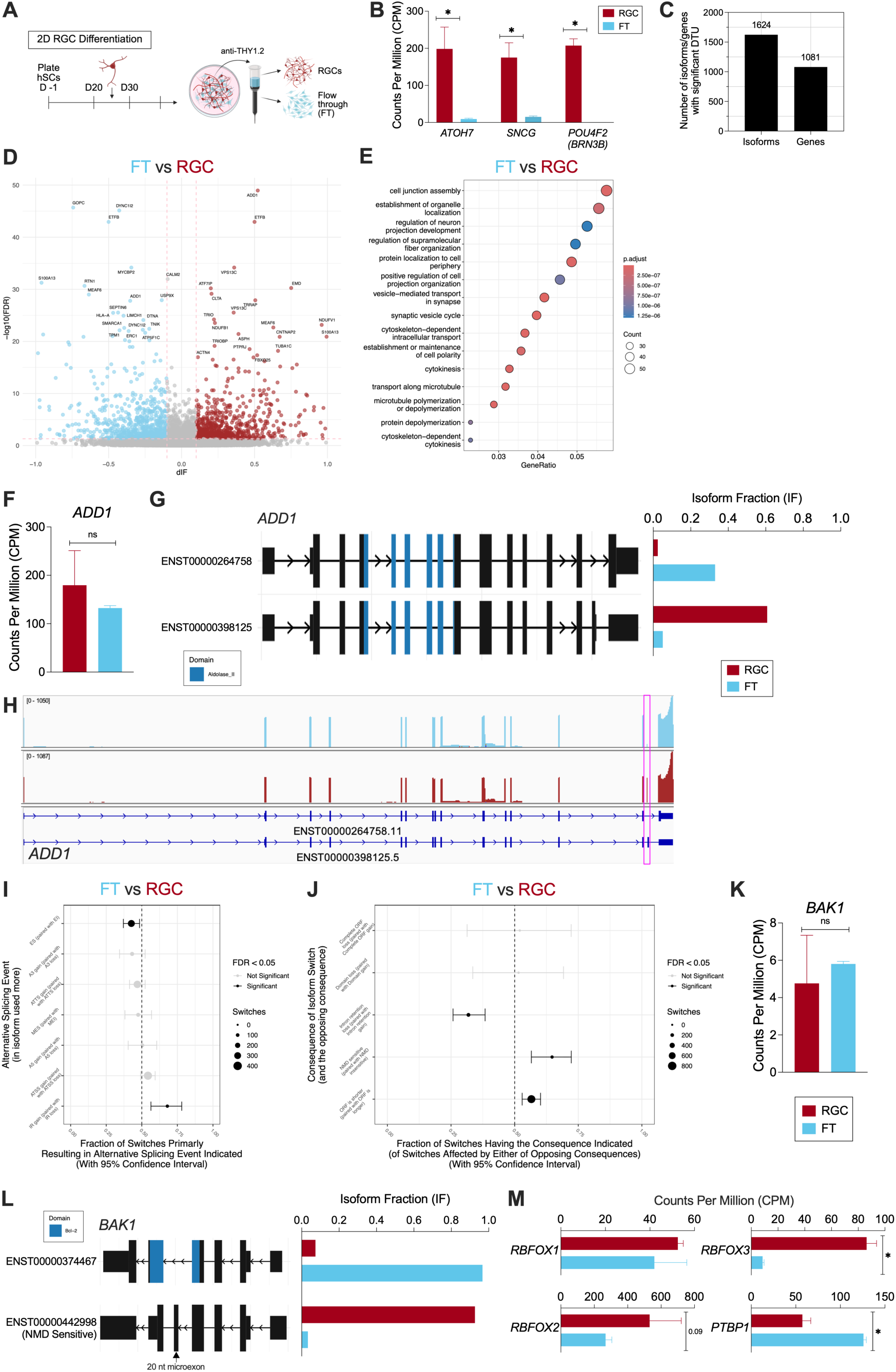
Cell type-specific alternative splicing profile of stem cell-derived retinal ganglion cells. **A)** Schematic of 2D differentiation and Magnetic-Activated Cell Sorting (MACS) purification of human stem cell-derived RGCs by Sluch et al. (55). Genome-engineered RGC reporter cells express tdTomato around Day 25 of 2D differentiation, which can be isolated using anti-THY1.2 magnetic beads to yield a pure population of RGCs and unbound flow-through for direct cDNA ONT sequencing. Created with BioRender.com. **B)** Gene-level expression (plotted as Counts Per Million, CPM) from long-read sequencing of RGC marker genes in purified RGCs and unbound flow-through (FT). Error bars indicate standard deviation. Asterisks denote gene is differentially expressed between groups (|log2FC| ≥ 1.0 & qval < 0.05). *ATOH7* – atonal bHLH transcription factor 7; *POU4F2* (*BRN3B*) – POU class 4 homeobox 2; *SNCG* – synuclein gamma. **C)** Number of isoforms and genes with significant DTU events (|dIF| ≥ 0.1 & qval < 0.05). **D)** Volcano plot highlighting differentially used transcripts between FT and RGCs. **E)** Top gene ontology (GO) biological process terms from genes with significant DTU comparing FT to RGCs. Heatmap indicates adjusted p-value for enrichment, and dot size represents the number of significant genes found within each GO term. **F)** *ADD1* (adducin 1) gene expression (plotted as counts per million, CPM). Error bars represent standard deviation. ns = not significant. **G)** Visualization of *ADD1* isoforms (left) with isoform usage of *ADD1* transcripts (plotted as isoform fraction, IF) (right). Colored exons denote conserved protein domain encoded within the mRNA transcript. Asterisks indicate significant DTU (|dIF| ≥ 0.1 & qval < 0.05). Aldolase_II – Aldolase II Superfamily, Class II Aldolase and Adducin head (N-terminal) domain. **H)** IGV browser showing reads preferentially mapping to an alternatively spliced cassette exon in the *ADD1* gene in RGCs (pink box). **I)** Types and enrichment of alternative splicing events found in genes with significant DTU. Dot size indicates number of significant DTU events. Red = significant skew of an alternative splicing event (qval < 0.05). **J)** Predicted consequences and enrichment of DTU events. Dot size indicates number of significant DTU events with predicted consequence. Red = significant skew of a consequence in the opposite direction (qval < 0.05). **K)** *BAK1* (BCL2 Antagonist/Killer 1) gene expression (plotted as CPM). Error bars represent standard deviation. ns = not significant. **L)** Visualization of *BAK1* isoforms (left), including a transcript containing a NMD-sensitive/poison microexon, with isoform usage of associated *BAK1* transcripts (plotted as IF). Bcl-2 – Apoptosis regulator proteins, Bcl-2 family. **M)** Gene expression of the RBFOX family of RNA-binding proteins and *PTBP1*, known regulators of *BAK1* poison exon splicing, in FT and RGCs (all plotted as CPM). Error bars represent standard deviation. Asterisks denote gene is differentially expressed between the two cell groups (|log2FC| ≥ 1.0 and qval < 0.05).

There were clear differences in alternative splicing patterns between RGCs and FT, particularly in cassette exon skipping/inclusion and intron retention (**Figure 7I**), which were coupled to differences in functional consequences like NMD sensitivity and ORF lengths (**Figure 7J**). One of the most significant DTU genes was *BAK1* (BCL2-antagonist/killer 1 protein). This pro-apoptotic gene has two mRNA isoforms, one of which includes a highly conserved microexon that functions as a “poison” exon when spliced in by generating a premature stop codon and subsequently triggering NMD (56). While *BAK1* gene expression levels did not significantly differ between RGCs and FT (**Figure 7K**), the NMD-sensitive transcript was overwhelmingly, and almost exclusively, the isoform used in RGCs, which was opposite to the FT samples where this isoform was nearly absent (**Figure 7L**). RT-PCR was used to confirm this alternative splicing pattern in both 2D RGCs and FT (**Figure S15**). Inclusion of this exon increased over time in the 3D retinal organoid samples, reflective of the higher proportion of terminally differentiated neurons expected in Stage 3 organoids (**Figure S15**).

Previous studies have shown that inclusion of this *BAK1* microexon is neural-specific and its splicing is regulated by RBPs like PTBP1 during neurodevelopment (56), which is also known to act in opposition with RBFOX family proteins to regulate microexon splicing transcripts in the brain (52). In our long-read data, we found that *RBFOX3* (with *RBFOX2* close but not meeting the adjusted p-value threshold for significance) was significantly upregulated in the RGCs compared to FT (**Figure 7M**). Conversely, *PTBP1* gene expression was much higher in the FT samples (**Figure 7M**), suggesting *BAK1* microexon splicing could be mediated through a similar mechanism in human stem cell-derived retinal neurons.

### Allele-specific expression (ASE) in stem cell-derived retinal cells

Long-read sequencing is advantageous for identifying ASE for multiple reasons. First, assigning reads to a specific allele/haplotype (phasing) is more accurate because the longer reads can span multiple sequence variants. Secondly, having longer reads is also advantageous when looking at other types of allele specificity, like allele-specific splicing events, that cannot be resolved as well, or at all, using short-read sequencing methods. These events can be located anywhere along the transcript and do not need to be nearby a variant to be detected. Using a read-based phasing approach, we performed ASE analysis on our direct cDNA ONT sequencing data. For this, phased heterozygous variants were used to assign the ONT reads to a haplotype, H1 or H2 (referred to as “haplotagging”). With this method, we could phase ∼25% of our ONT alignments across samples in all five retinal cell groups (Stage 1-3 organoids and 2D RGCs/FT), with no obvious bias in assignment to a particular haplotype (mean percentage of haplotagged reads across all samples = 13.4 ± 0.9% for H1 and 13.1 ± 0.8% for H2) (**Figure 8A**). We also did not observe any bias in the length of the reads successfully phased, since N50 read lengths for haplotagged reads were similar to the earlier reported, unphased read lengths (**Figure 1C, S14, S16**). Next, we performed DGE analysis to identify allelic expression imbalance between phased H1 and H2 reads within each retinal cell group (**Figure 8B, S16**). From these analyses, we found an enrichment of X chromosome genes that showed allele specificity (**Figure 8C, top**), which was apparent even when normalized to the number of genes encoded by each chromosome (calculated as a percentage; mean = 4.7% ± 1.3% for X chromosome ASE genes) (**Figure 8C, bottom**). While Stage 2 retinal organoids had the largest number of ASE genes (n= 119) of the five comparisons, there were no obvious differences in the distribution of these genes across chromosomes or the proportions of ASE imbalance towards H1 or H2 haplotypes compared to the other retinal cell groups (**Figure 8C**, **Figure S16**). Collectively, there was a large amount of overlap in ASE genes between all five retinal cell groups, with 27 genes showing ASE in all five comparisons (**Figure 8D**, **Table S9**).

**Figure 8.**
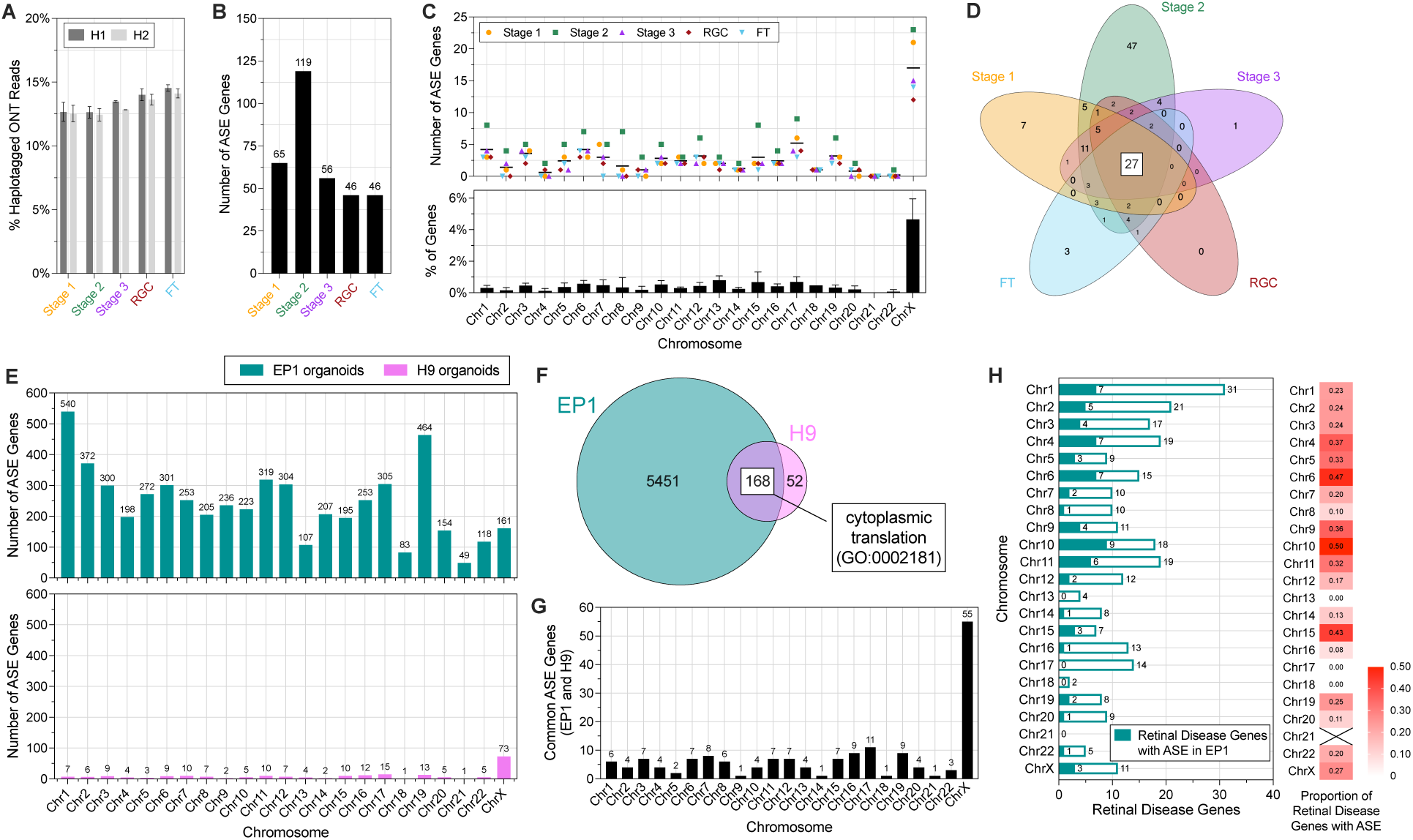
Allele-specific gene expression in stem cell-derived retinal cells. **A)** Percentage of haplotagged ONT reads for each cell group. Error bars indicate standard deviation between replicates. **B)** Number of genes with allele-specific expression (ASE) for each group (H1 vs H2). **C)** Number of ASE genes by chromosome (top) and mean percentage of ASE genes normalized to the number of genes encoded by each chromosome. Horizontal line denotes grand mean across all groups (top). Error bars denote standard deviation across all groups (bottom). **D)** Venn diagram showing the number of common and unique ASE genes between the different groups. **E)** Number of ASE genes by chromosome for EP1 (top) and H9 (bottom) background organoid samples. **F)** Venn diagram showing overlap of ASE genes between EP1 and H9 organoid samples. Genes related to the GO term “cytoplasmic translation” were enriched in the overlapping ASE genes. **G)** Number of ASE genes shared by EP1 and H9 analyses by chromosome. **H)** Number of retinal disease genes with ASE in EP1 samples by chromosome (left) and heatmap showing proportion of retinal disease genes with ASE (right).

### Differential ASE between pluripotent stem cell types

Despite finding many shared ASE genes amongst the grouped analyses, further examination of the individual samples revealed an unexpected clustering pattern of the phased gene counts by PCA. Rather than each sample/replicate being grouped by haplotype (H1 and H2), instead, we found that clustering was largely driven by the background, or cell origin, of the samples (**Figure S17**). Our long-read sequencing data on the differentiating retinal organoids included cells of two stem cell origins: induced pluripotent stem cells (iPSCs; EP1) and embryonic stem cells (ESCs; H9) (**Table S1**), while the 2D RGCs and FT replicates were only from an ESC-derived background, which strictly followed to the observed background-dependent clustering pattern. Several studies have found that DNA methylation and other histone marks can vary considerably between stem cell types (57, 58), which could impact ASE if also regulated in an allele-specific manner. To investigate differences in ASE between iPSC- and ESC-derived organoids, we grouped the organoid samples (since these included both stem cell types) by cell background (ie, EP1 or H9), rather than individual differentiation stages, and performed another DGE analysis comparing H1 and H2 haplotypes. This ASE analysis revealed that organoids derived from an EP1 background had vastly more ASE compared to H9-derived samples (**Figure 8E**). While we were able to phase almost identical percentages of H9 and EP1 reads to a haplotype (**Figure S17**), unlike in the earlier stage/cell type-specific ASE analyses where we found a disproportionate number of ASE genes were located on the X chromosome, ASE genes in EP1 were much more distributed across autosomes (**Figure 8E, top**). This differed from the ASE genes in H9-derived organoids, which followed the previously observed, X chromosome-enriched pattern for allelic imbalance (**Figure 8E, S17**). Noticeably, EP1 ASE genes were also more uniformly distributed across H1 and H2 haplotypes (**Figure S17**), which could be biologically significant or simply related to more genes reaching the established significance thresholds for ASE. We compared the ASE genes between EP1 and H9 samples and found that over 76% of H9 ASE genes where shared by the EP1 background samples (**Figure 8F**), 25% of these being X chromosome genes (**Figure 8G**). Of these 168 shared ASE genes, there was an overrepresentation of genes involved in “cytoplasmic translation” (GO: 0002181) but no other biological processes (**Figure 8F**), while GO analysis of EP1 ASE genes revealed an enrichment of autophagy and vesicle-related pathways (**Figure S18**).

Given the extent of allelic expression imbalance found in the EP1 samples, we were curious how many of the genes associated with retinal disease from our earlier analysis showed ASE in the EP1-derived organoids. Out of 274 retinal disease genes detected, 69 (25.2%) had significant ASE (**Figure 8H, Table S10**). These genes with ASE were not uniformly distributed across chromosomes, with a higher proportion of ASE retinal disease genes found on chromosomes 6, 10, and 15 (**Figure 8H**).

## Discussion

We could reliably capture the expected gene expression patterns of the retinal organoids in our long-read sequencing dataset, with high correlation to comparable short-read transcriptomic data, especially at earlier differentiation stages. Stage 3 long-read samples showed the lowest, but still good correlation to later organoid time points in the short-read dataset. Analysis of the genes driving this lower concordance found many retinal/photoreceptor genes were differentially expressed between the Stage 3 long-read and analogous short-read time points, which could be reflective of differences in the differentiation protocols used to produce the organoids for each of these studies. Notably, comparison of the two sequencing platforms also highlighted technical biases intrinsic to short-read data, as higher-expressing genes in the long-read samples were consistently longer had lower GC content (and vice versa). Given the limited number of long-read sequencing datasets for the retina, particularly human stem cell-derived retinal organoids, the observed gene-level concordance was overall encouraging and provided a convincing foundation to leverage our long reads for lesser defined isoform-level analysis in this *in vitro* model.

ONT sequencing produced reads that were ten-fold longer than what is typically obtained by conventional, short-read sequencing methods. These longer reads often spanned multiple exon-exon boundaries, allowing for reliable isoform quantification. The ability to capture greater isoform diversity (ie., more isoforms per gene) with the long reads was particularly apparent when directly compared to the transcripts detected in the short-read organoid dataset. Collectively, we identified thousands of isoform switching events through the generation of over 173 million direct cDNA ONT reads between 3D retinal organoid and 2D RGC cultures. These isoform switches frequently occurred in genes that were not differentially expressed. This was especially apparent in splicing-related genes that were largely expressed at high but unchanged levels across organoid developmental time points. DTU analysis of Parkinson’s disease brains also found substantial transcript usage changes non-DGE genes (59), further demonstrating the value of DTU analysis to uncover overlooked, and potentially important, temporally regulated transcriptomic changes during neuronal development.

Isoform switches can have substantial downstream impacts on cellular processes, resulting in altered protein-protein and/or protein-DNA interactions, potentially changing the function of a translated protein entirely (60, 61). One relevant case of this is basigin, or *BSG*, where the retina-specific isoform encodes for a protein containing an additional immunoglobulin extracellular domain that serves as the cell-surface receptor for rod-derived cone viability factor (62). IsoformSwitchAnalyzeR can predict large-scale patterns of functional consequences of isoform switches based on known protein annotations. While we could identify patterns of either enrichment or depletion for certain types of predicted consequences, these data still provide a rather superficial analysis of functional impact. Further exploration of the functional consequences of specific developmental splicing changes is intriguing and offers many exciting avenues for future studies.

We could detect multiple isoforms of genes associated with retinal disease in our long-read dataset. While many of these genes are predominantly expressed in photoreceptors and play critical roles in the development, function, and homeostasis of these cells, there was a subset of retinal disease genes that had significant DTU between younger and older organoid stages. Multiple isoforms of *CRB1*, including a retina-specific transcript, were previously identified through PacBio sequencing of both mouse and adult human retinal tissue (18). While this study reported previously unannotated *CRB1* transcripts, our ONT reads mapped to nine *CRB1* isoforms in the GENCODE reference annotation. Of these, three had significant DTU in the retinal organoids and expression of ENST00000367397, which closely resembles the retina-specific variant in Ray et al.’s study, progressively increased in our organoid samples over time (**Figure 4D, S12**).

Neural-specific splicing patterns were found in our long-read sequencing data of 2D, enriched RGCs. One such hallmark of neuronal cells is the high inclusion of microexons, a subset of highly conserved, alternatively spliced cassette exons with lengths of 3-27 nucleotides (corresponding to 1-9 amino acids) (63). We found several transcripts containing microexons were preferentially used the 2D RGCs, such as the NMD-sensitive *BAK1* isoform involved in the suppression of apoptosis following neurogenesis (56). We also identified DTU genes associated with neurological diseases, like *ADD1*. A recent study found that *ADD1* splice variants are differentially expressed between cortical progenitor cells and post-mitotic neurons (64), which matches the *ADD1* isoform usage patterns we detected in the RGCs and unbound cells. Moreover, studies have shown that optic nerve degeneration occurs in adducin knockout mice (64, 65), further highlighting its importance in RGCs. In the case of both *BAK1* and *ADD1* splicing, PTBP1 acts as a suppressor of the neuronal transcript (56, 64), which is likely regulated in a similar manner in our stem cell-derived neurons.

Our initial analysis looking at allelic expression imbalance across five retinal cell groups found an enrichment of allele specificity in X chromosome genes. X chromosome inactivation is a well-recognized and necessary process for silencing one copy of the chromosome in female cells to equalize gene dosage, which can occur randomly in cells of female origin, which were used in this study. However, this phenomenon can erode in prolonged human stem cell cultures, and to varying extents in iPSC and ESC lines(66, 67). While we do not know the degree of X inactivation erosion in our stem cell cultures, one of the common ASE genes we found in our analyses, phosphoglycerate kinase 1 (*PGK1*), is considered a “textbook” example of X inactivation where expression of either the maternal or paternal allele is temporally regulated during development (68). The most remarkable finding from our ASE analysis seemed to be the magnitude of ASE genes in iPSC background cells compared to samples of embryonic origin. Characterization of allele specificity in human ESCs and iPSCs has been largely in the context of X inactivation (69), with little known about the ASE status for autosomal genes. It has been shown that iPSCs retain their donor ASE profiles, which can vary between individual donors/lines, a potentially confounding factor contributing to iPSC variability (70). Furthermore, epigenetic patterns can also significantly vary between pluripotent cell types, especially at earlier passages (57). Taken together, this initial analysis offers compelling directions for future allele-specific analysis of both gene and isoform-level expression, as well as allele-specific modifications on both the DNA and RNA (e.g., methylation) that could regulate this process. Further validation of these ASE patterns and experiments to understand the extent of ASE variability amongst different stem cell lines and between donor backgrounds seems warranted and highlights a potentially important factor to consider when using human stem cell models.

Despite being a limited analysis in terms of sample size for the iPSC-derived retinal organoids, it is still worth noting that we found a substantial number of genes implicated in retinal disease with significant allelic expression imbalance, which were not equally distributed across chromosomes. Interestingly, two of the chromosomes with the highest proportion of ASE retinal disease genes, chromosomes 6 and 10, both encode genes/loci associated with age-related macular degeneration (AMD). While AMD is a multifactorial disease with associated risk factors such as demographics (e.g., sex, aging, ethnicity) and environment (e.g., sunlight, cigarette smoke), several genetic variants have also been linked to the disease (71, 72). One of the strongest genetic risk loci for AMD is the *ARMS2/HTRA1* region (73–75) located on chromosome 10, which did exhibit ASE in the EP1 samples (**Table S11**). Notably, Li et al. found evidence of ASE in several complement pathway genes that have known associations with AMD (76–79), including *CFB* (located on chromosome 6), *CFH*, and *C3*, in their transcriptomic studies of post-mortem human eye tissues (80). Although none of these genes showed ASE in the EP1 samples with our significance cut-offs, it is still notable that *CFB* expression significantly differed between haplotypes and its adjusted p-value fell just under our threshold for significance (log2FC = −5.34, adjusted p-value = 0.055) (**Table S11**). Understanding the allele specificity of AMD genetic risk factors, as well as the Mendelian retinal disease genes, in retinal stem cell models may be an important factor to consider when investigating phenotype heterogeneity and disease pathogenesis, especially as the popularity of *in vitro* disease modeling with iPSCs continues to rise.

Our bulk long-read sequencing study spanning retinal organoid differentiation time points offers a comprehensive investigation of the dynamic changes in isoform usage during human retinal organoid development and will hopefully serve as a valuable resource to the vision research field, as well as the broader neuroscience community. Nevertheless, long-read sequencing technologies continue to evolve rapidly, with major improvements in both chemistry and analysis pipelines released on a frequent basis. While we were able to achieve over 99% mapping of our reads to the reference genome, direct cDNA sequencing is known to generate a higher proportion of chimeric reads compared to other ONT protocols, such as cDNA PCR-based ONT sequencing, due to library preparation artifacts (81). In addition, while long-read sequencing has opened new opportunities for transcriptome-wide isoform discovery, benchmarking studies have not yet converged on which computational tools provide the most accurate detection of novel isoforms. In our analysis, we observed substantial differences between results from Bambu and IsoQuant, reflecting the variability that currently exists across algorithms. To mitigate this uncertainty, we adopted a consensus-based strategy, focusing on isoforms that were consistently identified by both tools. As long-read chemistries improve transcript coverage and new analysis methods continue to mature, we anticipate that future work will achieve greater reliability and sensitivity in the discovery of novel isoforms. In addition, our long-read sequencing data captured many non-coding RNA biotypes in the developing organoids, including 22.7% lncRNAs (**Figure S1E**). While our analyses largely focused on isoform usage changes in protein-coding genes, examination of the non-coding retinal transcriptome is another area of RNA biology worthy of in-depth study.

Future studies leveraging updated kits and basecalling models are likely to provide even more comprehensive and reliable insights into retinal organoid transcriptome complexity. Although we identified temporal isoform usage events across organoid development, whether these isoform switches reflect changes in cell type composition of the differentiating organoids or differential usage within the same retinal cell type (or both), this cannot be resolved with bulk long-read transcriptomics alone. To this point, while our isoform-level analysis of the 2D stem cell-derived RGC cultures allowed us to examine alternative splicing events for a single retinal cell type, it is still difficult to make direct comparisons to the organoids since the differentiation methods (3D organoids versus 2D RGCs) are inherently different. ONT’s platform is now capable of sequencing of full-length cDNA directly from single cell gene expression libraries for identification of isoforms at individual cell resolution. Nanopore sequencing was recently combined with short-read RNA-sequencing for single cell isoform analysis of mouse retinal tissue (20). Future single cell long-read studies using human-derived retinal organoids will further enhance our understanding of not only temporal splicing changes but also of the landscapes of isoform usage across different cell types in the developing retina.

### Conclusions

We conducted one of the first long-read RNA sequencing studies on human stem cell-derived retinal organoids to investigate the dynamics of isoform usage during retinogenesis. Our findings demonstrate the independence of overall gene expression levels and relative isoform specificity, emphasizing the importance of understanding isoform diversity and the alternative mRNA splicing landscapes that coincide with the long and intricate processes of retinal cell development. Together, these results deepen our understanding of post-transcriptional regulation of the central nervous system and provide meaningful insights into human stem cell-derived organoids in hopes of expanding their future utility in understanding molecular mechanisms of human embryonic development and disease.

## Methods

### Stem cell lines and cell culture

H9 (WA09) ESCs modified with BRN3B-tdTomato (55) and CRX-tdTomato (82), EP1 stem cells (83), and EP1 cells modified with BRN3B-tdTomato (84) were used for this study (**Table S1**). All research involving human stem cells was conducted in accordance with the International Society for Stem Cell Research Guidelines with approval from the Institutional Stem Cell Research Oversight Committee at Johns Hopkins University. Stem cells were cultured on growth factor reduced Matrigel^©^ Growth Factor Reduced Basement Membrane Matrix, LDEV-free (Corning) in mTeSR™ Plus medium (STEMCELL Technologies) and maintained in a 37°C incubator at 5% CO_2_. For passaging, cells were detached using Accutase™ solution (Sigma-Aldrich) and seeded in mTeSR™ Plus medium containing 5 μM (–)-Blebbistatin (Sigma-Aldrich). All cell lines were passaged at least two times before differentiation.

### Retinal organoid differentiation

Retinal organoids were generated using a protocol adapted from Cowan et al. (27). In brief, stem cells were dissociated using Accutase™ solution (Sigma-Aldrich) for seeding into wells (pre-treated with Anti-Adherence Rinsing Solution (STEMCELL Technologies)) of AggreWell™ 800 microwell culture plates (STEMCELL Technologies) in mTeSR™ Plus medium containing 10 μM (–)-Blebbistatin and transitioned to neural induction medium (NIM: DMEM/F12 GlutaMAX™ supplement (Gibco), 1X N-2 Supplement (Gibco), 1X MEM Non-Essential Amino Acids Solution (NEAA, Gibco), and 2 μg/mL heparin sodium salt from porcine intestinal mucosa (Sigma-Aldrich) for embryoid body (EB) formation. After one week of culture, the EBs were adhered to Matrigel^©^ Growth Factor Reduced Basement Membrane Matrix-coated 6-well plates, followed by the switch to a 3:1 ratio of DMEM, high glucose, GlutaMAX™ Supplement, pyruvate (Gibco) and Ham’s F-12 Nutrient Mix, GlutaMAX™ Supplement (Gibco) containing 1X B-27™ Supplement, minus vitamin A (Gibco), 1X MEM NEAA, and 1X Antibiotic-Antimycotic (Gibco). After three weeks of adherent culture, the optic vesicles (OVs) were detached and transferred to ultra-low attachment 6-well plates for further differentiation and maintenance of retinal organoids in suspension for up to 30 weeks as previously described (27).

### 2D retinal ganglion cell (RGC) differentiation and purification

H9 ESCs modified with BRN3B-tdTomato (also engineered to express a unique cell surface protein, THY1.2) were differentiated into RGCs using a previously described protocol (55). Stem cells were seeded and differentiated as a monolayer. Cells were differentiated to approximately Day 40, then purified by magnetic-activated cell sorting (MACS) with CD90.2 (THY1.2) MicroBeads (Miltenyi Biotec) as previously described (55).

### Total RNA extraction

RNA was extracted using TRIzol™ Reagent (Invitrogen) (85). Four to six retinal organoids were pooled at three stages of differentiation: Week 6/Day 45 (Stage 1), Week 14/Day 100 (Stage 2), and Week 30/Day 210 (Stage 3). Pooling was performed in duplicate (Stage 1 and Stage 3) or triplicate (Stage 2) using independent stem cell lines as biological replicates (**Table S1**). Each organoid stage included replicates from both H9 and EP1 backgrounds. For MACS-purified 2D RGCs and unbound column flow-through (FT) cells, RNA was extracted from approximately 1 million cells for each sample in triplicate from two independent differentiations. Purified RNA was quantified using a Qubit 3.0 Fluorometer (Thermo Fisher Scientific) with an RNA High Sensitivity Assay Kit (Thermo Fisher Scientific), and RNA Integrity Numbers (RINs) were determined using an Agilent 2100 Bioanalyzer and RNA 6000 Pico Kit (Agilent).

### Direct cDNA sequencing

Full-length, PCR-free cDNA was prepared using a reverse transcriptase and strand-switching method and the Ligation Sequencing Kit V14 from ONT. Approximately 1 μg of total RNA was used as input for cDNA preparation. cDNA libraries were quantified using a Qubit dsDNA High Sensitivity Assay Kit (Thermo Fisher Scientific), and library length was measured using a Bioanalyzer system with an Agilent High Sensitivity DNA Kit. Each cDNA library was loaded onto a single PromethION R10.4.1 flow cell (ONT) following the manufacturer’s protocol and sequenced on a PromethION 24 device (ONT) using MinKNOW’s default run settings.

### Read alignment and transcript quantification

Basecalling was performed in real-time using Guppy version 6.5.7 (https://nanoporetech.com/software/other/guppy). FASTQ files were filtered based on Phred quality score (Q-score ≥ 10) using NanoFilt (86) version 2.8.0. Minimap2 (87) version 2.1 was used to align filtered reads to the GENCODE human release 46 (GRCh38.p14) reference annotation (88). Per-read junction distribution was calculated from these alignments. Known and novel transcript identification and quantification was performed using Bambu (89) version 3.7.0 and IsoQuant (90) version 3.6.0 on primary mapped alignments with mapping quality (MAPQ ≥ 30). Out of the 1,118 novel isoforms discovered by Bambu, 496 were also identified by IsoQuant. Only these novel isoforms were used for downstream filtering. For transcript filtering, known and novel isoforms were only retained if they arose from protein-coding genes. Counts-based filtering was performed with edgeR (91) version 4.0.14 using *filterByExpr* with *min.count* set to ten (genes) or two (isoforms). Transcript filtering is outlined in **Figure S2**.

### Differential analyses

All downstream analyses were performed in R version 4.3.2. Isoform counts matrices were input into IsoformSwitchAnalyzeR (92) version 2.2.0 for Differential Transcript Usage (DTU) analysis using the DexSeq method (36) with the default counts-filtering disabled. Transcripts were identified as differentially used if the differential isoform fraction |dIF| ≥ 0.1 and adjusted p-value (or q-value) ≤ 0.05 using the false discovery rate (FDR) method (93). edgeR was used to perform Differential Gene Expression (DGE) and Differential Transcript Expression (DTE) without running expression-based filtering since this was already done in the prior step. Genes and transcripts were considered differentially expressed if |log2FC| ≥ 1.0 and adjusted p-value ≤ 0.05 using FDR.

### Short-read RNA-sequencing data processing

FASTQ files from independent, publicly available bulk short-read RNA-sequencing data of human stem cell-derived retinal organoids (31) were downloaded from NCBI BioProject #PRJNA754196. This dataset included 28 samples across seven time points: D0, D10, D25, D65, D100, D180, and D280. Reads were quantified using Salmon (35) version 1.10.1 against the GENCODE release-46 primary-assembly transcriptome with a plain (decoy-free) salmon index (salmon index -k 31, 251,955 targets) using salmon quant −l A --validateMappings --gcBias --seqBias. Per-sample quants were aggregated with tximport (94) version 1.37.2 to a gene matrix (62,266 genes × 28) and, with txOut=TRUE, a transcript matrix (251,955 × 28). For quantifying exon-exon junctions, the short reads were genome-aligned with STAR (95) version 2.7.8a (release-46 primary assembly, sjdbOverhang = 149), and for each uniquely mapped read (MAPQ = 255) the number of junctions crossed was counted as the number of N operations in its CIGAR string, restricted to multi-exon protein-coding genes.

### Comparison of long- and short-read RNA sequencing data

To compare the two platforms without the quantifier as a confounder, the long-read organoid samples were re-quantified with salmon in *alignment mode* (salmon quant -a --noErrorModel) on the existing minimap2 transcriptome BAMs (-ax map-ont), against the same release-46 FASTA used for the short reads. These were transformed with tximport to a gene matrix (62,700 genes × 7) and a transcript matrix (252,835 × 7). Short- and long-read gene matrices were CPM-normalized (edgeR) and subset to the protein-coding genes detected in common between the two platforms (19,993 genes; the mitochondrial rRNA gene ENSG00000210082 was excluded as an outlier). Spearman correlation was computed between every long- and short-read sample. DGE was performed using edgeR at three matched developmental stages (**Table S3**). Normalization factors were computed by TMM (calcNormFactors on the long-read samples and short-read samples independently), then combined. A post-normalization CPM ≥ 1 filter was applied within both platforms, and the long-read matrix was subset to protein-coding genes to match biotype scope. A gene was called differentially expressed if |log2FC| ≥ 1.0 & FDR < 0.05.

For each differentially expressed genes, median transcript length and GC content across its GENCODE isoforms were taken from the release-46 transcript metadata, and the distributions of these genes were compared for each organoid differentiation stage. Exon-exon junctions spanned per read were used to compare read lengths using the aligned data for each platform (long-read counts from minimap2 and short-read counts from STAR). To compare transcript-level matrices from both platforms, a detection filter of ≥ 1 estimated count in ≥ 1 sample defined the detected isoform set. To compare the number of isoforms per gene from each platform, transcript matrices were subset to protein-coding isoforms and filtered with edgeR filterByExpr (*min.count* = 2), followed by TMM for the CPM matrices (the deposited count matrices are the raw filtered counts). Isoforms per gene were counted on the filtered matrices and binned per sample (1–10, and 10+).

### Splicing enrichment and functional consequence analyses

For analyzing specific types of splicing events between isoforms in a significant switch, the *extractSplicingEnrichment* function was run in IsoformSwitchAnalyzeR. For analysis of the functional consequences of isoform switches, the coding potential of isoforms was predicted using CPC2 version 1.0.1 (96) (Python 3 version) and IsoformSwitchAnalyzeR was used to generate the corresponding protein sequences. These sequences were scanned for protein family domains using Pfam Scan (97), and SignalP (98) version 6.0 was used to predict the presence of signal peptides. Integration of the results from these external tools was done with the *IsoformSwitchAnalysisPart2* function to summarize functional consequences of the isoforms, including intron retention, coding potential, open reading frame (ORF) sequence similarity, nonsense-mediated decay (NMD) status, and identified signal peptides and domains (and isotypes). The *extractSplicingSummary* and *extractConsequenceSummary* functions were run to acquire the number of significant isoform switches with the above functional consequences.

### Overrepresentation analysis of retinal disease-associated genes

GraphPad Prism version 11 was used to calculate odds ratios and perform one-sided Chi-square tests (at a 95% confidence interval) to measure the extent of enrichment of multi-isoform genes and DTU frequency in retinal disease-associated genes relative to all genes included in the transcriptome-wide analyses.

### Allele-specific expression (ASE) analysis

Publicly available whole genome sequencing data from human H9 ESCs was obtained from the NCBI SRA database (SRX347299), aligned to the GENCODE 46 release using Bowtie2 (99), and merged into a single bam file. Quality control was performed Genome Analysis Toolkit (100) best practices pipeline (101) including deduplication of reads with *MarkDuplicates* and base recalibration with *BaseRecalibrator* and *ApplyBQSR*. Only primary mapped reads with a Mapping Quality (MAPQ) score greater than 20 were retained, which was done using samtools (102). The filtered bam file was used to create a variant call format (VCF) file using GATK’s *HaplotypeCaller*. Single nucleotide variants (SNVs) and indels in the resulting file were filtered using GATK best practices filters (**Figure S19**). A phased VCF was generated by passing the filtered VCF and long-read RNA sequencing bam files through WhatsHap (103, 104). This read-based phasing tool requires reads to cover at least two heterozygous variants; reads that covered only a single heterozygous variant (singletons), or no variants, were unphased. On average (across all chromosomes), 88.2 ± 2.1% of the heterozygous variant calls were phased, with a median phase set/block size of 3 variants (mean = 11.03 ± 1.94 variants) (**Figure S20**). Next, the phased VCF was used to assign the long reads to a haplotype using *whatshap haplotag.* Reads that did not cover any heterozygous variants were excluded. To help mitigate reference-mapping bias, the –reference flag was used both in whatshap phase and whatshap haplotag commands so that reads were locally re-aligned against both the reference and alternative allele; if both alleles were equal, the read was labeled as “unknown” (105). Haplotagged reads were subset to perform gene-level quantification using Bambu (89). Allele-resolved counts were arranged as a protein-coding gene matrix, where each column represented one haplotype (H1/H2) of a sample at a given differentiation stage. After removing genes with zero counts across all columns, expressed genes were retained with edgeR::filterByExpr (min.count = 3, grouping columns by haplotype × stage). The threshold was lowered from the default of ten because allele-specific quantification assigns only reads over phased heterozygous sites to a haplotype (∼25% of alignments). Counts were TMM-normalized, and genes with |log2FC| ≥ 1.0 & FDR < 0.05 were considered differentially expressed (allele-specific).

### Data visualization

The *prcomp* function from baseR was used to perform PCA on log2-transformed transcripts per million (TPM) and plotted using ggplot2 (106) version 3.4.4. Transcript expression heatmaps were plotted using ComplexHeatmap (107) version 2.18.0. Gene ontology (GO) analysis and visualization was performed using the *enrichGO* and *dotplot* functions from clusterProfiler (108) version 4.13.0. UpSetR (109) version 1.4.0 was used to create upset plots for DTU genes and transcripts across the different retinal organoid stages. The Integrative Genomics Viewer (IGV) user interface (110) version 2.16.2 was used to visualize exon usage in select genes. The Venn diagram to visualize shared ASE genes between groups was made with InteractiVenn (111). Additional figure generation was done with GraphPad Prism version 11.

## Declarations

### Ethics approval and consent to participate

Not applicable.

### Consent for publication

Not applicable.

### Availability of data and materials

Raw and processed data are available in GenBank with BioProject PRJNA1171814 and as a R/Bioconductor ExperimentHub package https://github.com/sparthib/HumanRetinaLRSData. The code for this project is publicly available through GitHub at https://github.com/sparthib/retina_lrs and is permanently archived on Zenodo under DOI: 10.5281/zenodo.21815470.

### Authors’ contributions

C.J.K. and D.J.Z. conceived and designed the study. Funding was acquired by C.J.K., S.C.H., and D.J.Z, and W.T., S.C.H., and D.J.Z. provided the resources for this study. Experiments were performed by C.J.K., Y.C., X.S., X.C., E.Y., and S.C. The code was implemented by S.P., and S.C.H., and C.J.K., S.P., and S.C.H. analyzed the data. C.J.K. and S.P. prepared the original draft of the manuscript with review, editing, and approval by all authors.

## Acknowledgements

The authors would like to thank David Gamm at the University of Wisconsin-Madison for providing the H9-CRX-tdT cell line, and the Johns Hopkins University & Medicine Genetics Resource Core Facility for performing some of the ONT sequencing runs included in this study. The authors also thank the maintainers of the Joint High Performance Computing Exchange (JHPCE) compute cluster at Johns Hopkins Bloomberg School of Public Health for providing essential computing resources. The authors also thank the Smith 3^rd^ Floor Writing Accountability Group for support during the drafting and editing of the manuscript.

## Supplemental Methods

### RT-PCR/qRT-PCR validation

Validation of the ONT sequencing data was done with independent differentiations of retinal organoids and 2D RGCs using the same protocols described above. Total RNA was isolated from individual retinal organoids and 2D purified RGCs and FT cells (n = ∼500K cells each) using an miRNeasy micro kit (Qiagen) according to the kit protocol. The purified RNA was reverse transcribed using the High-Capacity cDNA Reverse Transcriptase Kit (Applied Biosystems) following the manufacturer’s protocol. qRT-PCR primers were designed to amplify specific mRNA isoforms. Primer amplification efficiencies were assessed by a 5-fold dilution series standard curve, and post-amplification melt curve analysis was done to verify primer specificity. qRT-PCR was set up in technical triplicate using SSoAdvanced Universal SYBR Green Supermix (Bio-Rad Laboratories) following the instruction manual and run on a CFX384 Real-Time System (Bio-Rad Laboratories). qRT-PCR data analysis was performed using the Delta C_t_ method with the geometric mean C_t_ values of two housekeeping genes (*GAPDH* and *ACTB*). For RT-PCR, primers were designed to amplify a region that included an alternatively spliced cassette exon. RT-PCR was performed using Phusion High-Fidelity PCR Master Mix with HF Buffer (Thermo Scientific) following the user guide recommendations for setting up and running the PCR reactions in a T100 Thermal Cycler (Bio-Rad Laboratories). Annealing temperatures for each RT-PCR primer set were optimized by gradient PCR. Densitometric analysis was performed using ImageStudio™ Lite (LICOR bio) to quantify the proportion of exon inclusion and exclusion. All primer sequences used in this study are included in **Table S7**.

**Figure S1.**
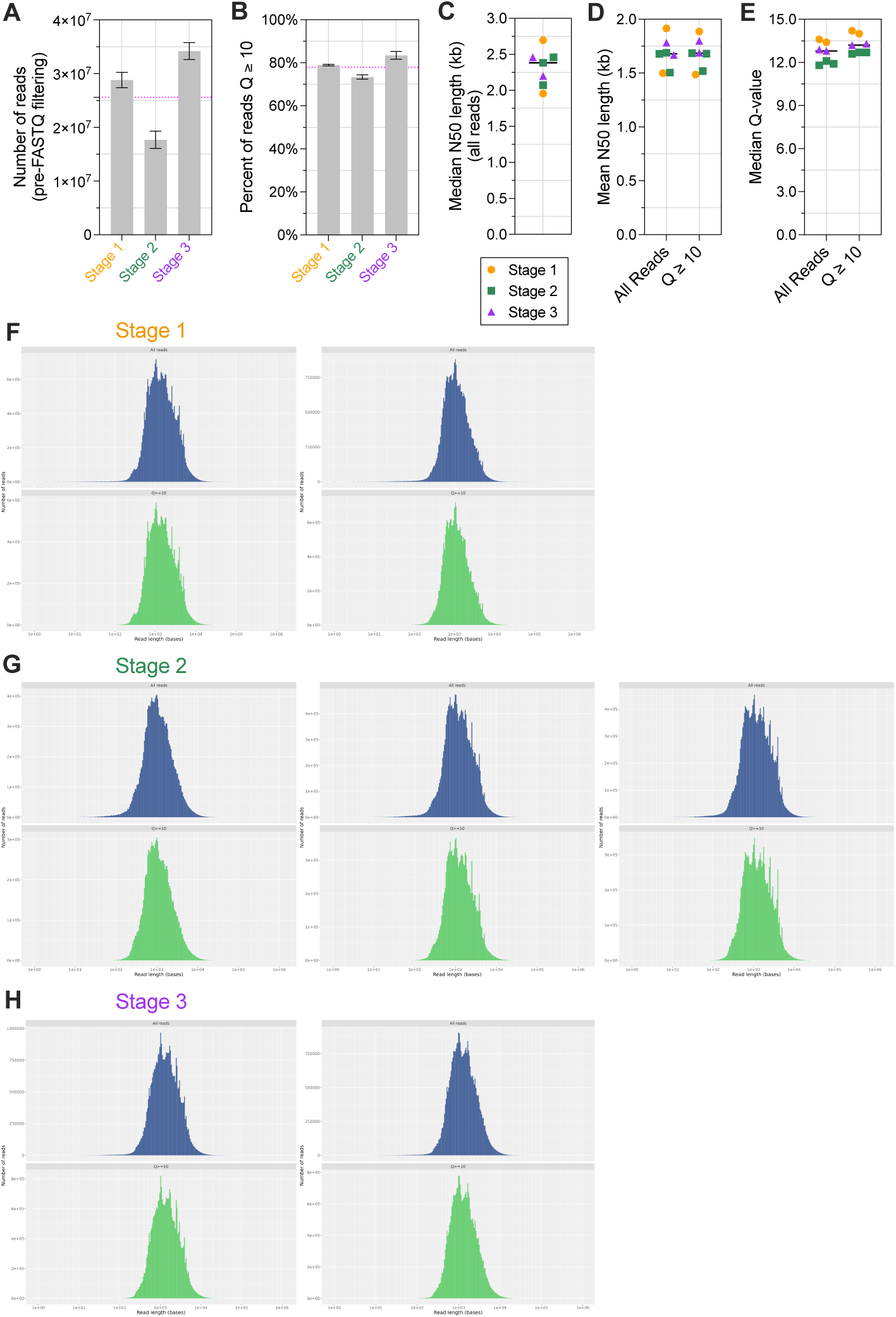
ONT read statistics for all organoid samples. **A)** Total number of reads pre-FASTQ filtering. **B)** Percent of reads (pre-FASTQ filtering) with Q-score ≥ 10. For A and B, error bars denote standard deviation and dashed pink lines indicate average across all organoid samples. **C)** Median N50 length for all reads. **D)** Mean N50 length of all reads and reads with a Q-score ≥ 10. **E)** Median Q-score for all reads and reads with Q ≥ 10. For D-F, black line denotes grand mean across all samples. **F-H)** Histograms of ONT read lengths for all reads (top) and reads with a Q-score ≥ 10 (bottom) for **F)** Stage 1, **G)** Stage 2, and **H)** Stage 3 replicate samples.

**Figure S2.**
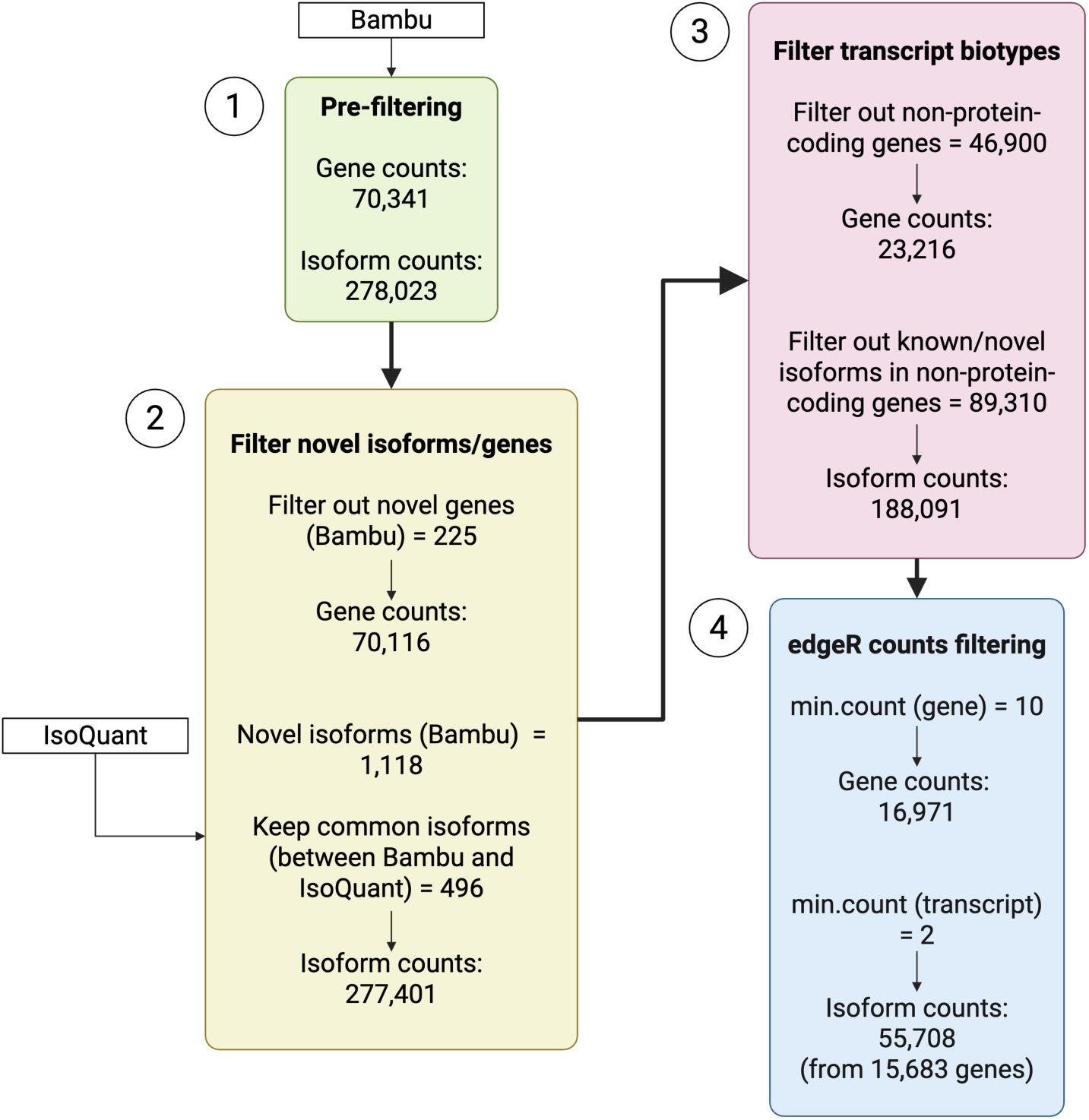
Flowchart for isoform/gene counts filtering of retinal organoid ONT sequencing samples. Created in BioRender.com.

**Figure S3.**
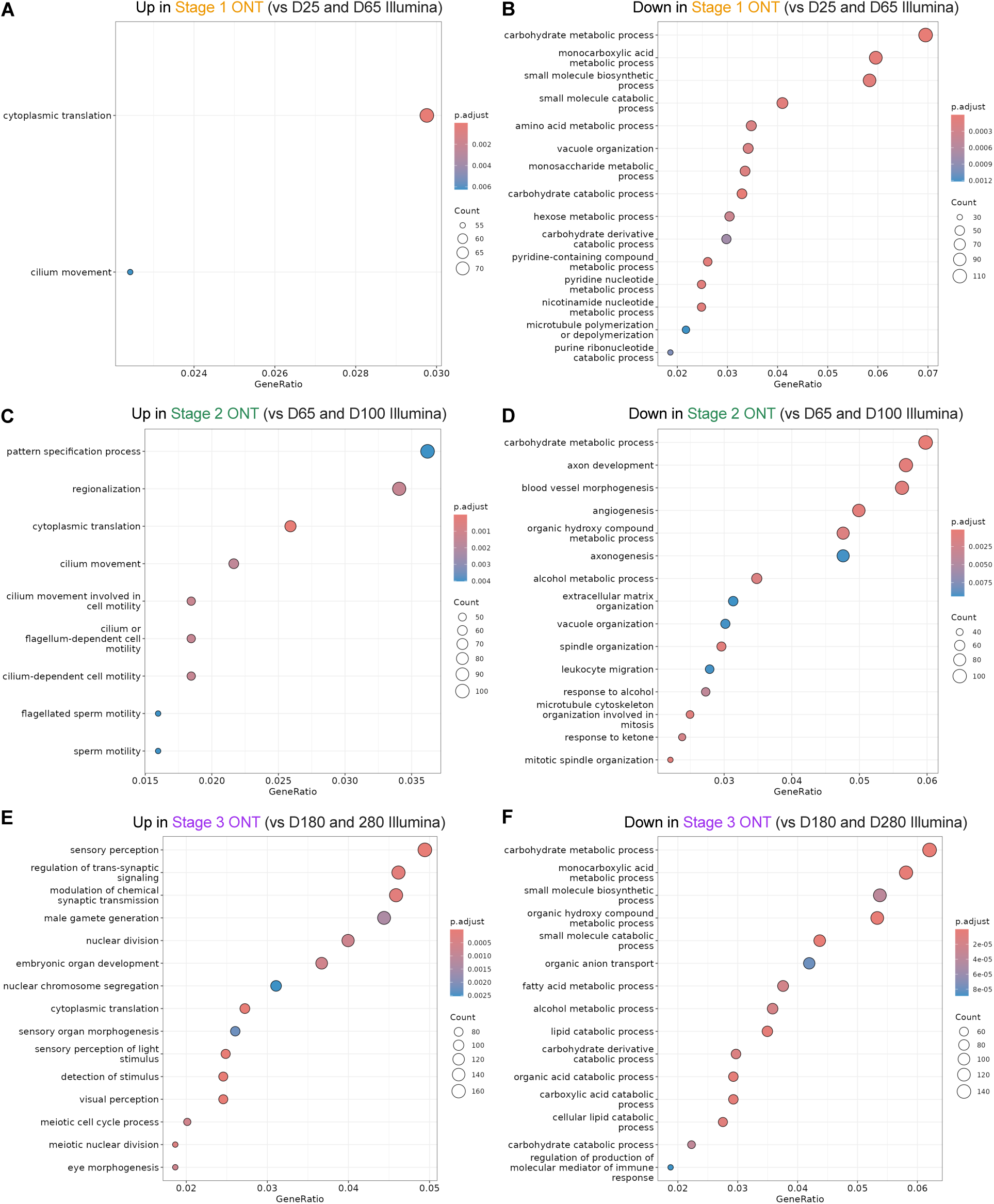
GO analysis of differentially expressed genes in organoids across ONT and Illumina platforms. **A-B)** Stage 1 ONT vs D25 and D65 Illumina, **C-D)** Stage 2 ONT vs D65 and D100 Illumina, and **E-F)** Stage 3 vs D180 and D280 Illumina time points.

**Figure S4.**
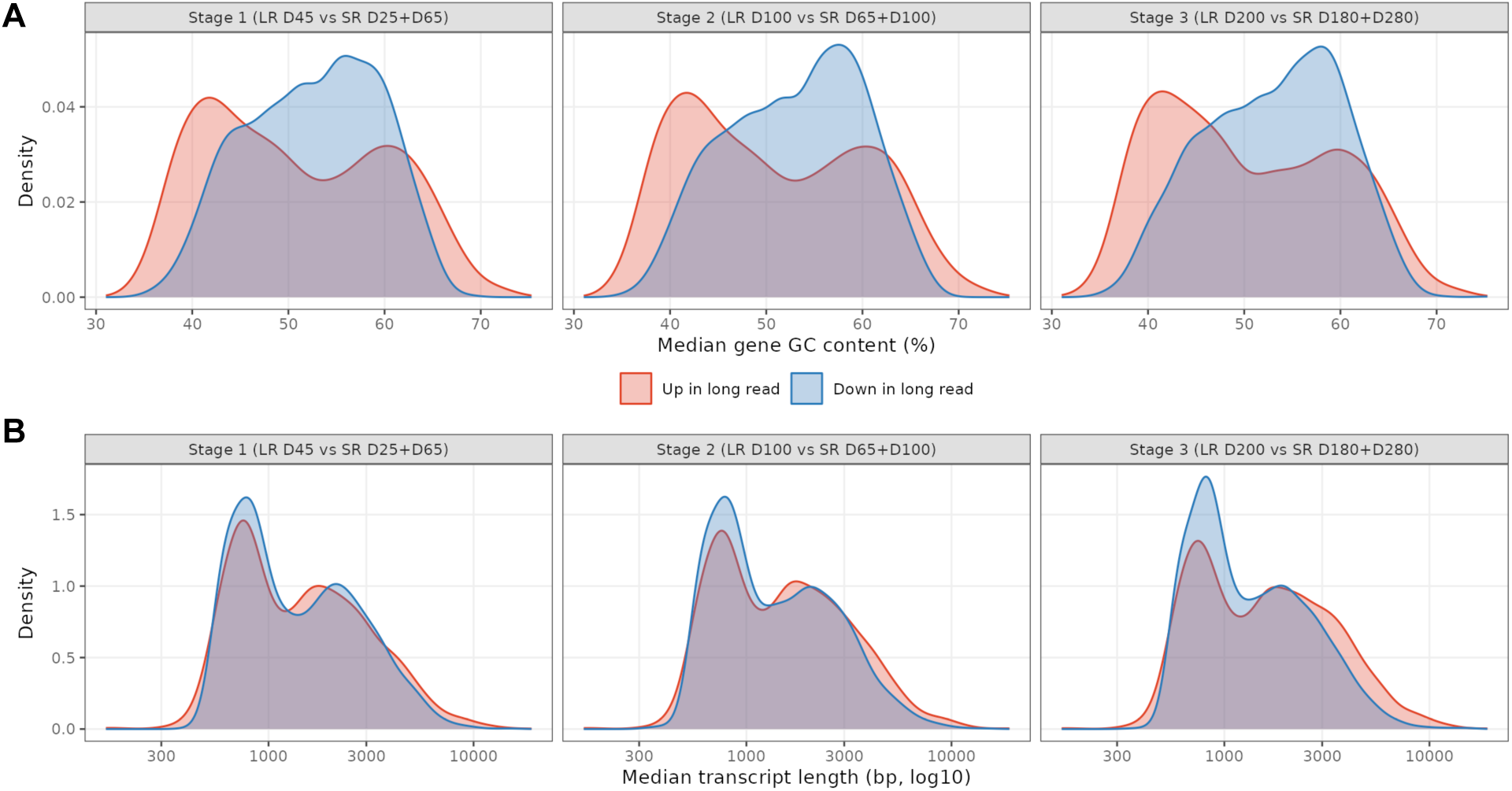
Technical comparison of long-read and short-read platforms. **A)** Median GC content (as percentage, %) for differentially expressed genes in retinal organoids across long- and short-read platforms. **B)** Median transcript lengths (as log10 base pairs, bp) for differentially expressed genes across ONT and Illumina platforms. LR – long-read (ONT); SR – short-read (Illumina).

**Figure S5.**
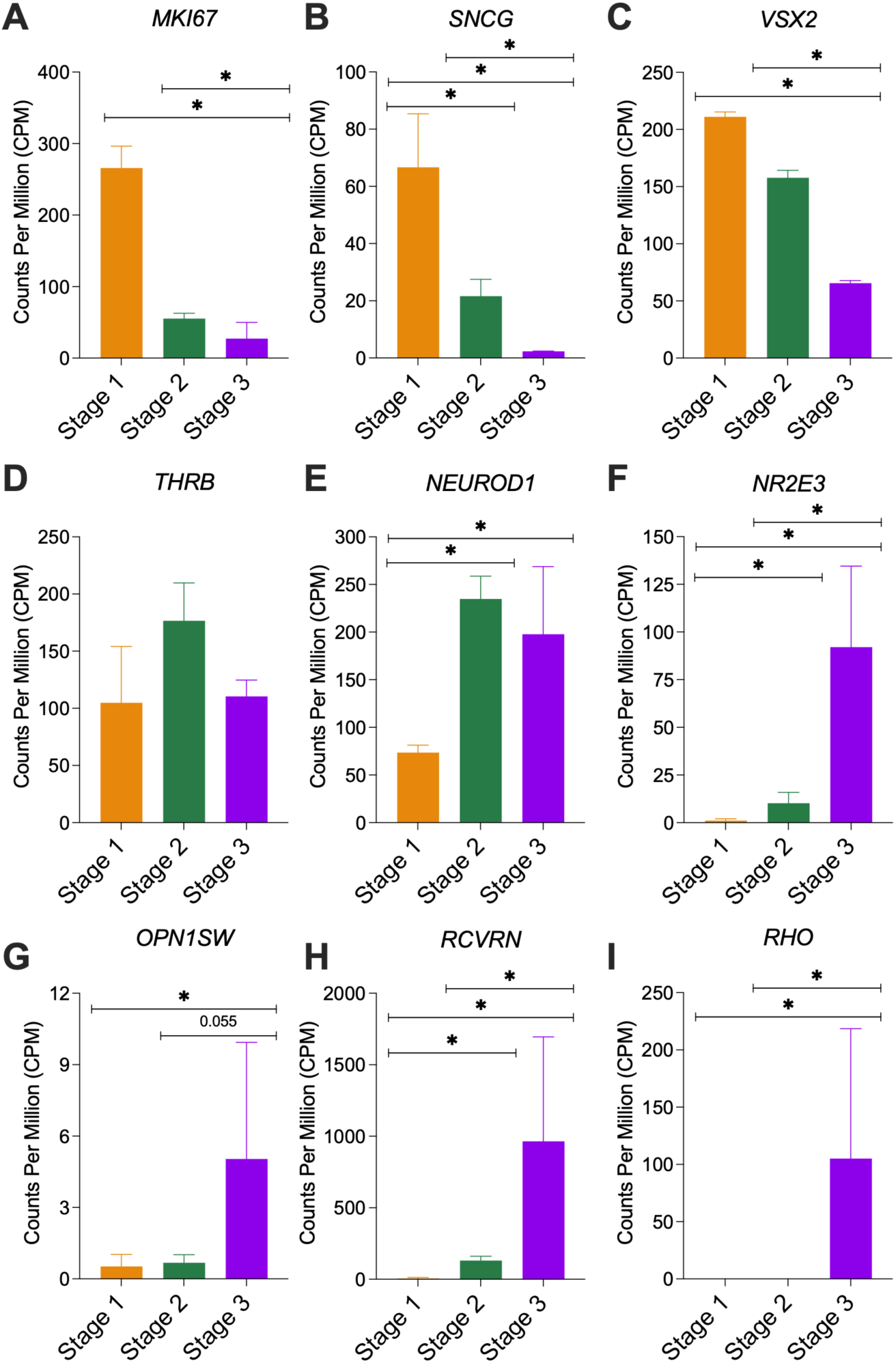
Gene expression (as Counts per Million, CPM) of additional retinal organoid/cell type markers. **A)** *MKI67* (mitotic cells) **B)** *SNCG*, (RGCs) **C)** *VSX2* (retinal progenitors), **D)** *THRB* (cone precursors), **E)** *NEUROD1* (neurogenic progenitors/neurons), **F)** *NR2E3* (rod photoreceptors), **G)** *OPN1SW* (cone photoreceptors), **H)** *RCVRN* (rod photoreceptors), and **I)** *RHO* (rod photoreceptors). Error bars show standard deviation. * indicates differential gene expression.

**Figure S6.**
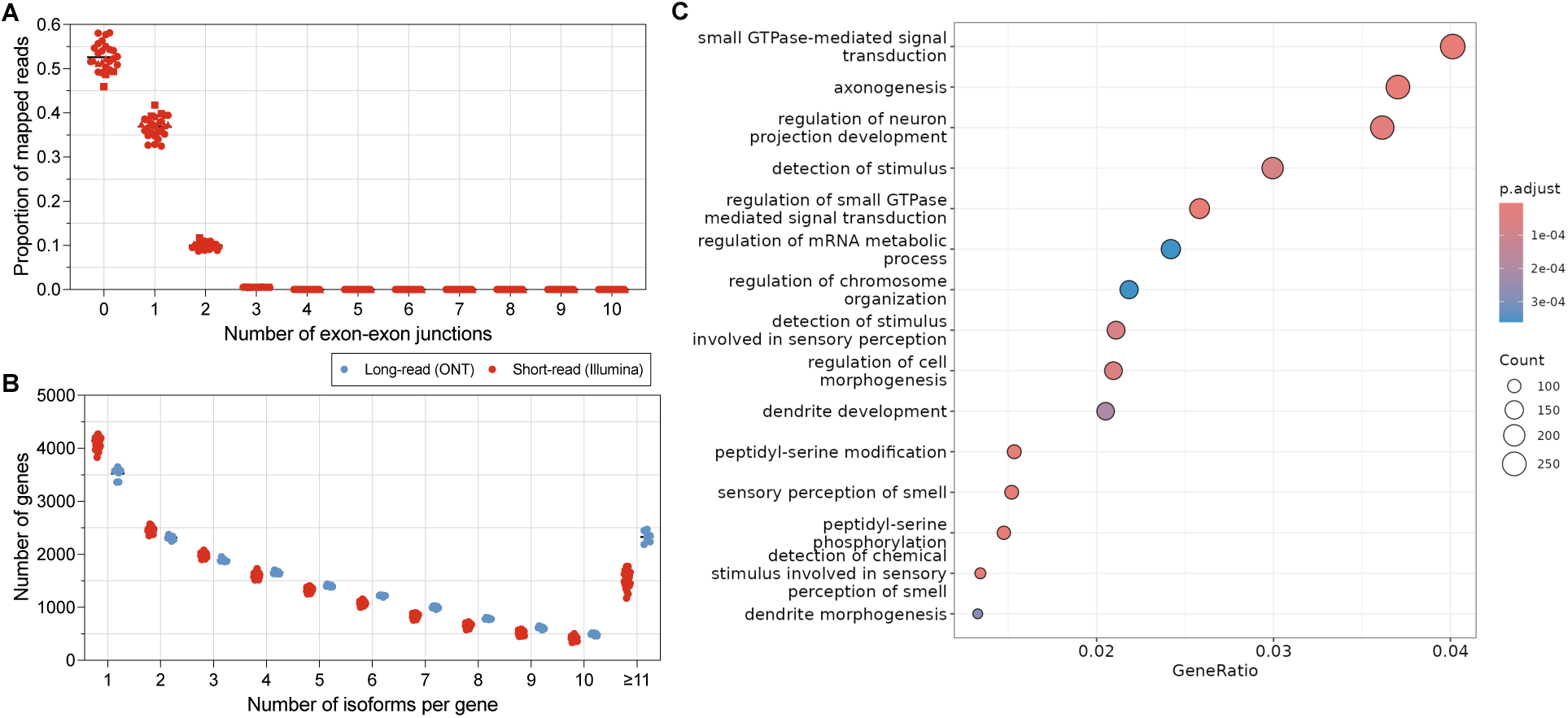
Isoform-level analysis comparing long- and short-read RNA sequencing data for human retinal organoids. **A)** Number of exon-exon junctions spanned by the Illumina reads. **B)** Histogram showing the number of isoforms detected per gene (quantified using salmon for both platforms) in organoid samples across both long- and short-read platforms. **C)** GO analysis for genes with isoforms unique to long-read samples.

**Figure S7.**
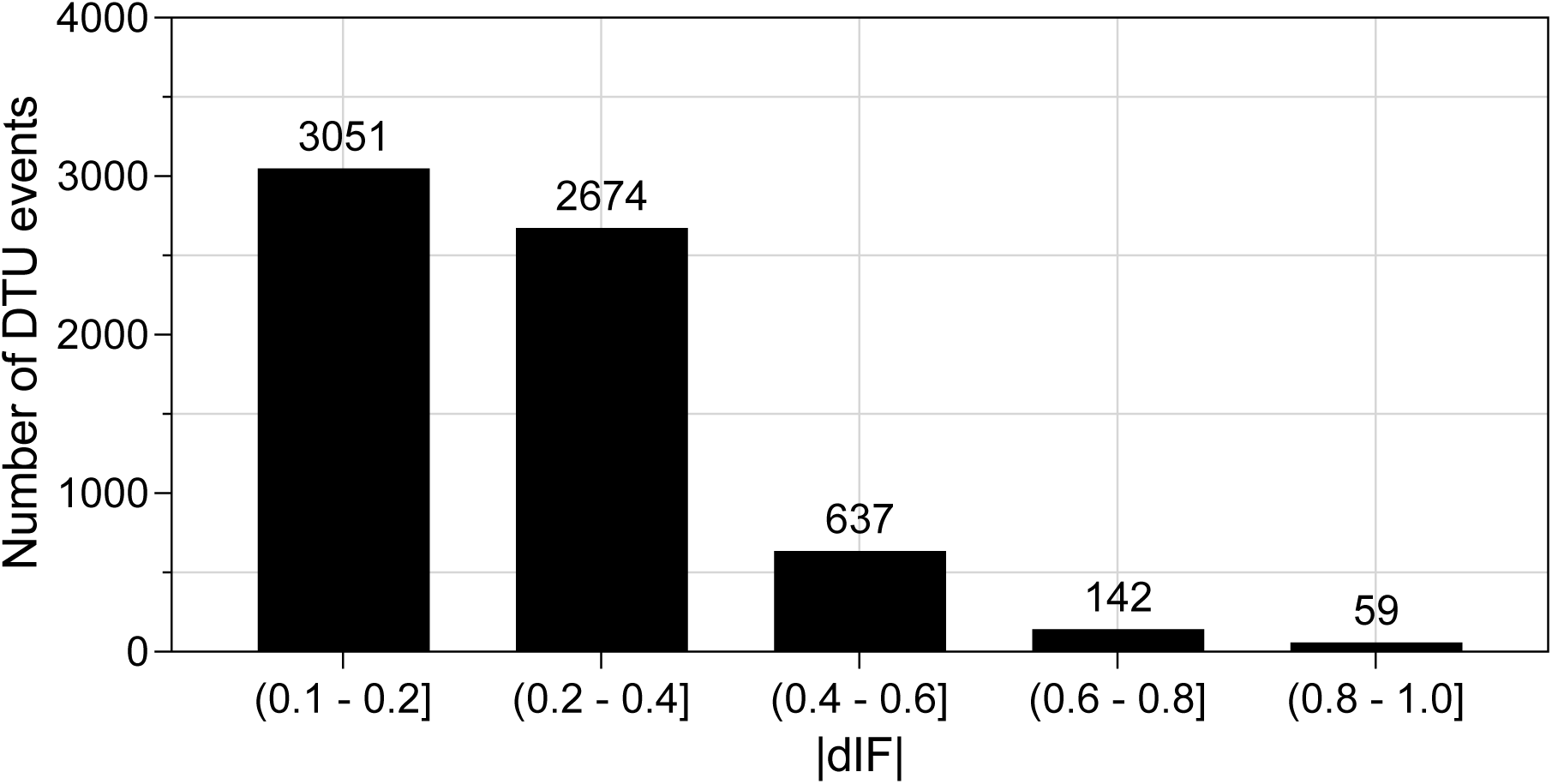
Differential isoform fraction (dIF) for significant DTU events. Histogram of dIF values for significant DTU events (|dIF| ≥ 0.1 & qval < 0.05) across all three comparisons in retinal organoids. The average (mean) dIF was 0.250.

**Figure S8.**
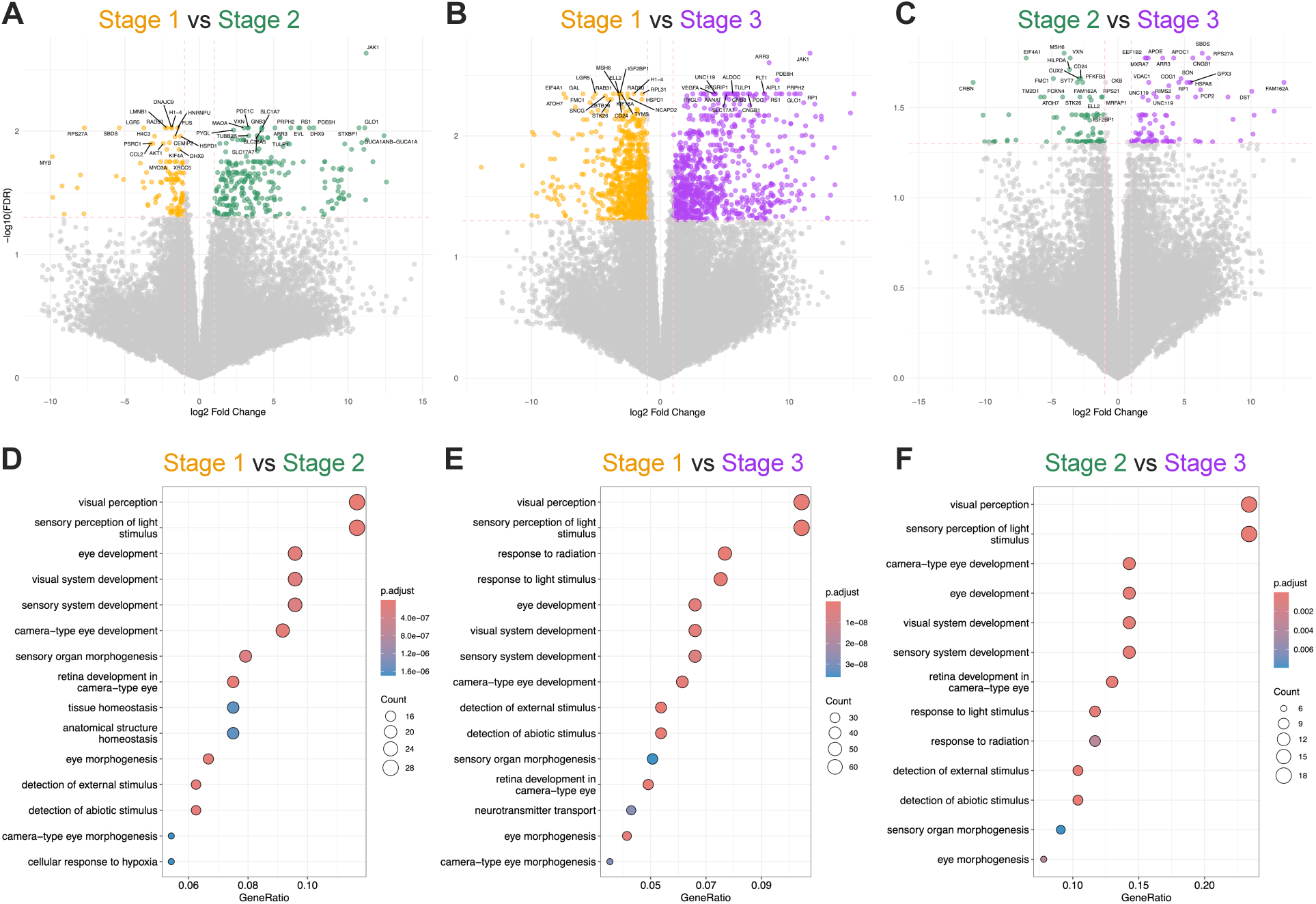
Differential transcript expression (DTE) between retinal organoid time points. **A-C)** Volcano plots showing DTE transcripts for **A)** Stage 1 vs Stage 2, **B)** Stage 1 vs Stage 3, and **C)** Stage 2 vs Stage 3 retinal organoid samples. **D-F)** GO analysis of genes with differential transcript expression in **D)** Stage 1 vs Stage 2, **E)** Stage 1 vs Stage 3, and **F)** Stage 2 vs Stage 3 retinal organoids.

**Figure S9.**
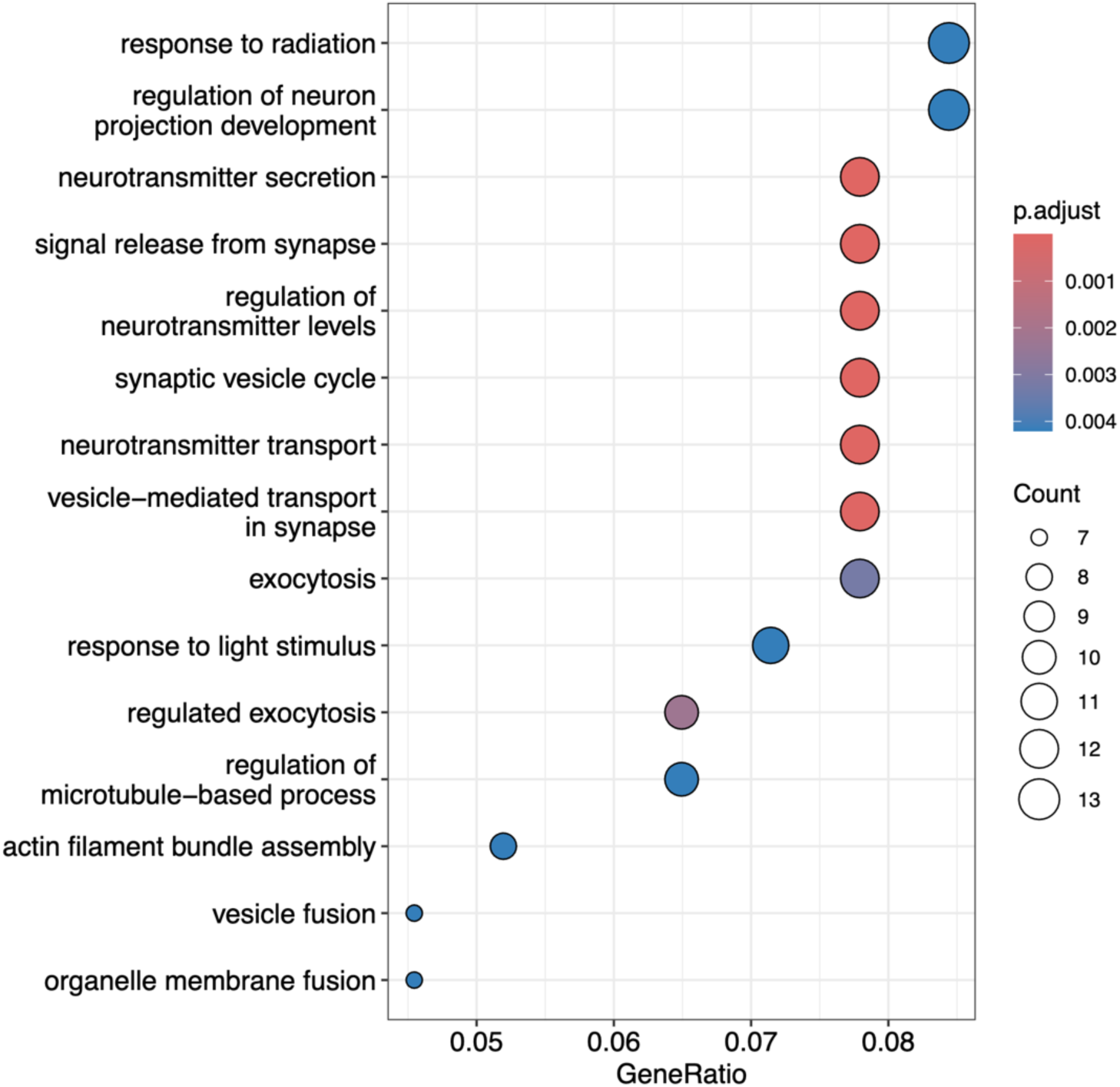
Gene Ontology (GO) analysis of common genes with isoform switches.

**Figure S10.**
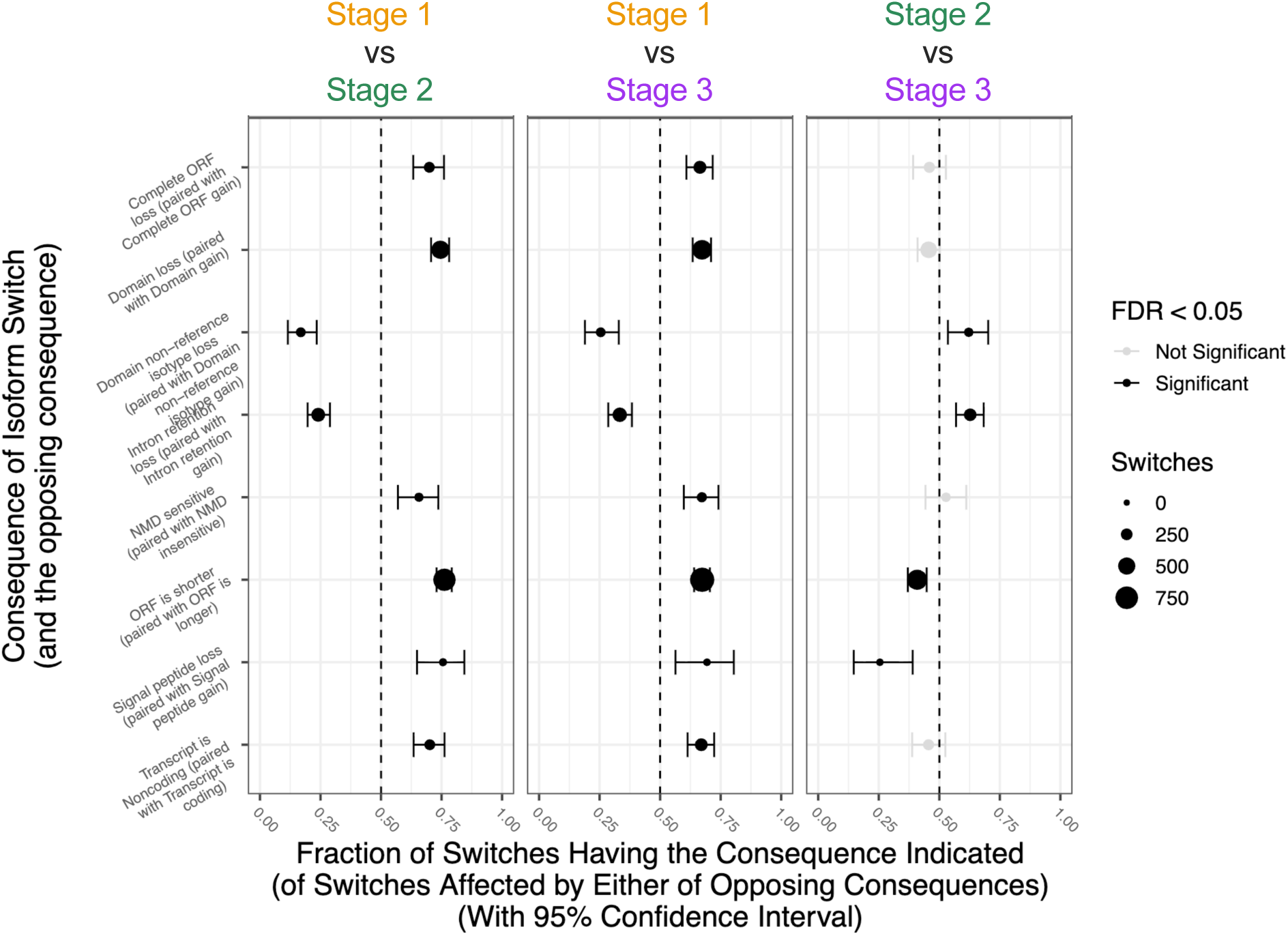
Functional consequences of isoform switches in retinal organoids. Size of dot indicates number of significant DTU events. Black denotes significant shift (FDR < 0.05) in either enrichment or depletion of a functional consequence.

**Figure S11.**
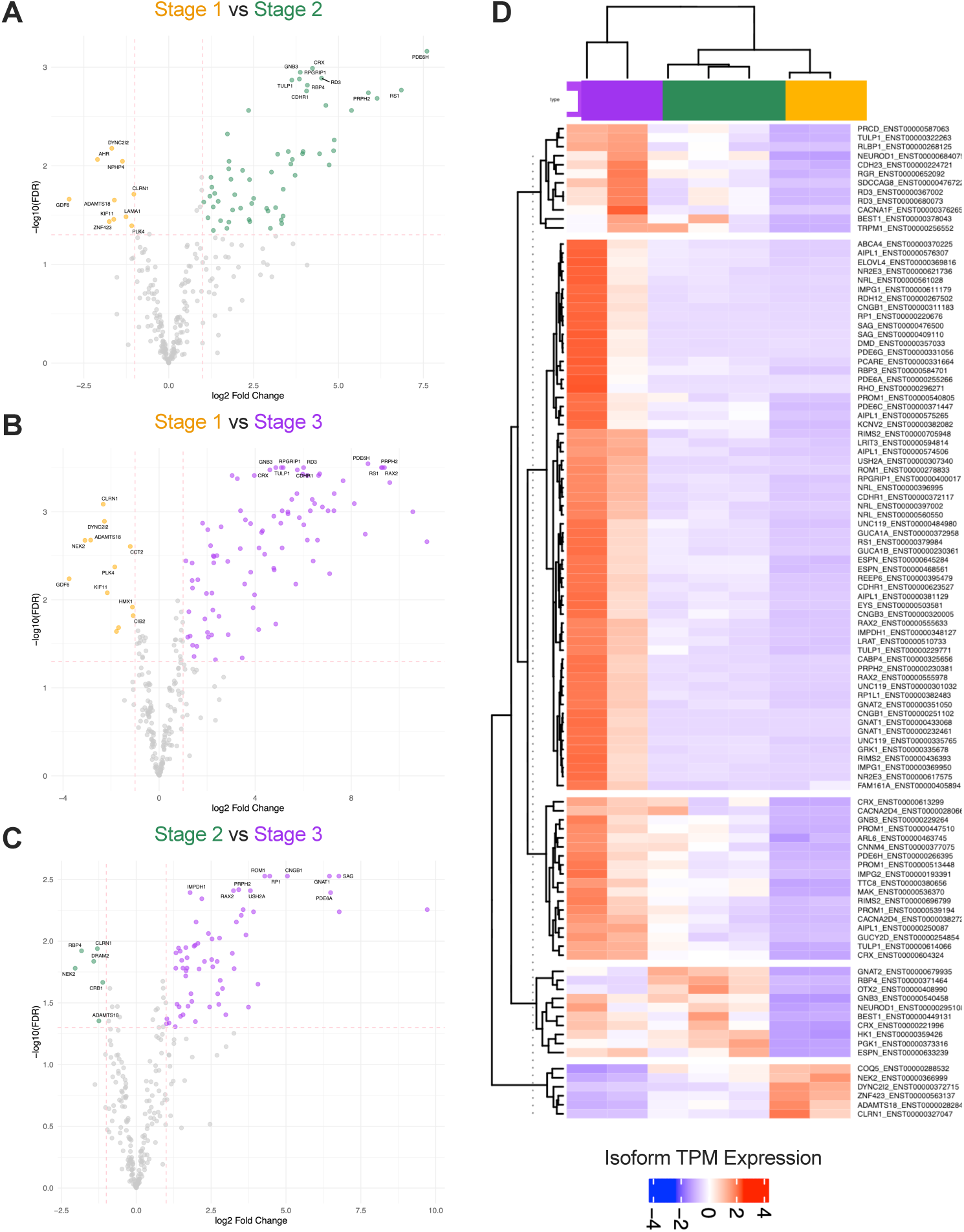
Differential gene and isoform expression in retinal disease genes. **A-C)** Volcano plots showing differentially expressed retinal disease genes in **A)** Stage 1 vs Stage 2, **B)** Stage 1 vs Stage 3, and **C)** Stage 2 vs Stage 3 retinal organoids. **D)** Heatmap of differentially expressed isoforms of retinal disease genes. Heatmap colors denote scaled isoform transcripts per million (TPM) expression.

**Figure S12.**
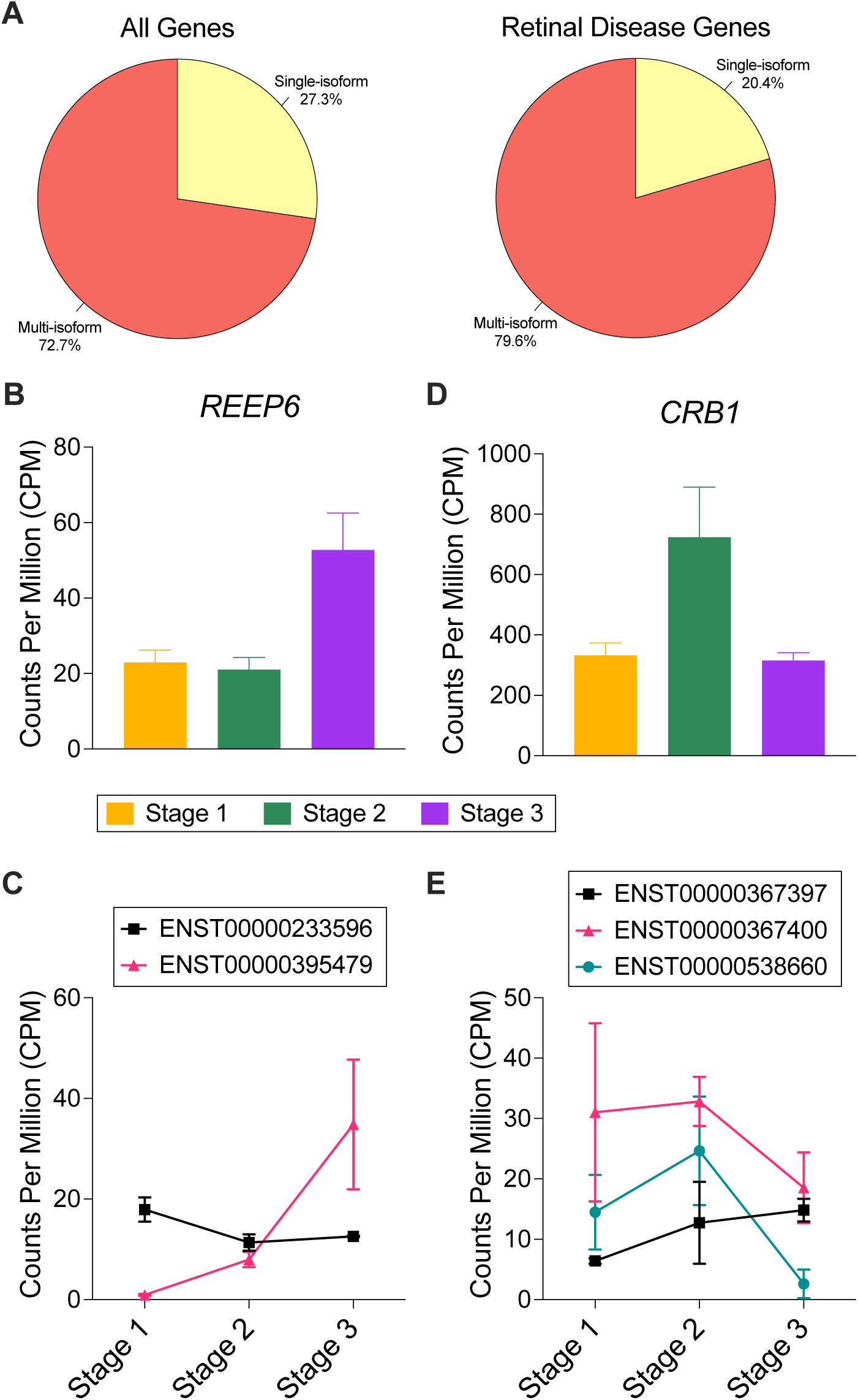
Isoforms of retinal disease-associated genes. **A)** Proportion of single-isoform and multi-isoform genes of all genes analyzed (left) and the subset of retinal disease genes (right). **B-C)** *REEP6* **B)** gene and **C)** isoform expression in differentiating retinal organoids. **D-E)** *CRB1* **D)** gene and **E)** isoform expression in organoids. For B-E, error bars denote standard deviation.

**Figure S13.**
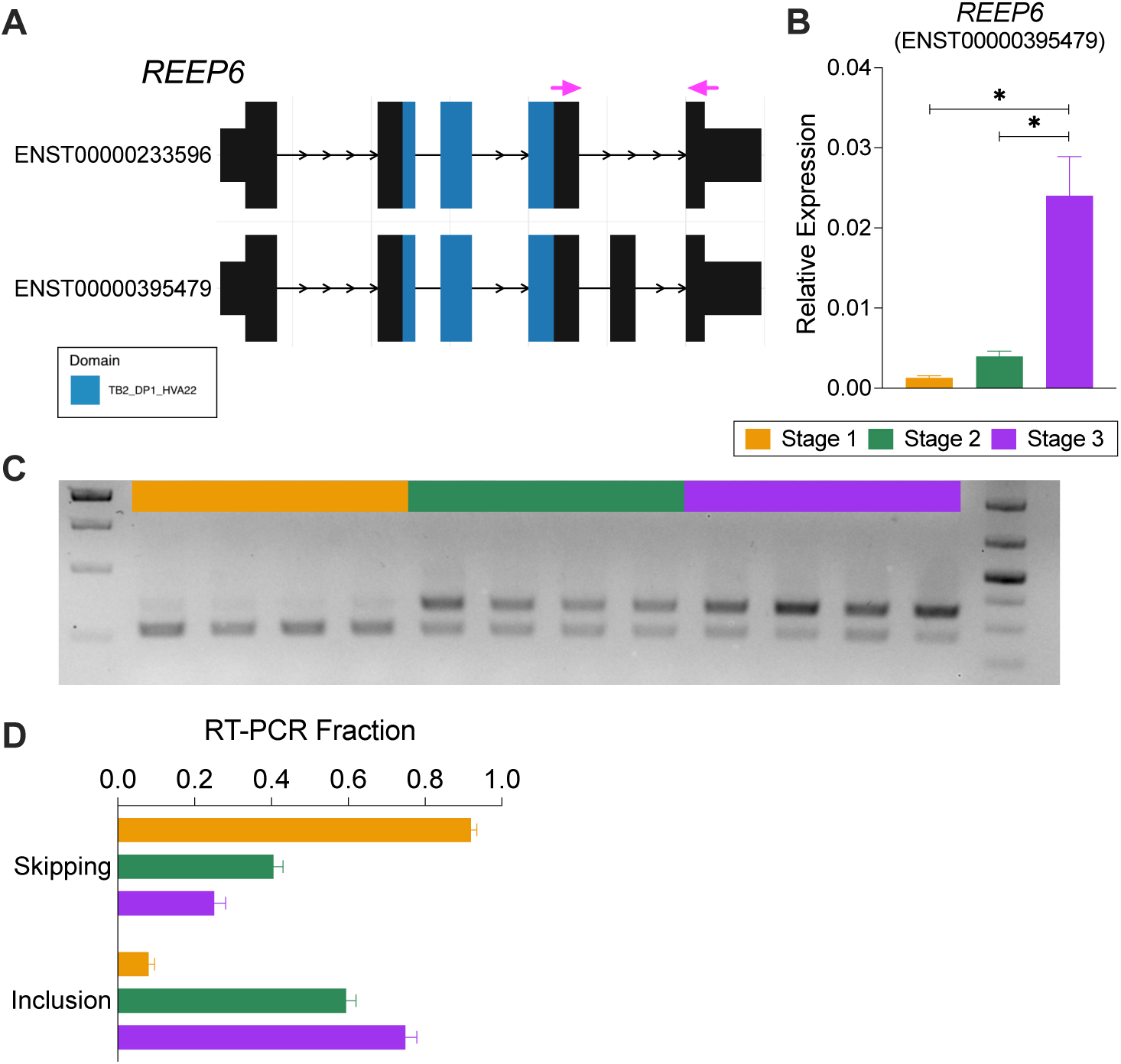
Validation of *REEP6* splicing in retinal organoids. **A)** Schematic of *REEP6* isoforms. Pink arrows denote RT-PCR primers flanking the alternatively spliced cassette exon. **B)** qRT-PCR using primers specific to the ENST00000395479 variant. Error bars show SEM, n=8 replicate organoids per group. * indicates significance between groups (One-Way ANOVA). **C)** Agarose gel image of *REEP6* RT-PCR of the alternatively spliced cassette exon in retinal organoids. **D)** Quantification of *REEP6* RT-PCR cassette exon skipping and inclusion. Error bars indicate SEM.

**Figure S14.**
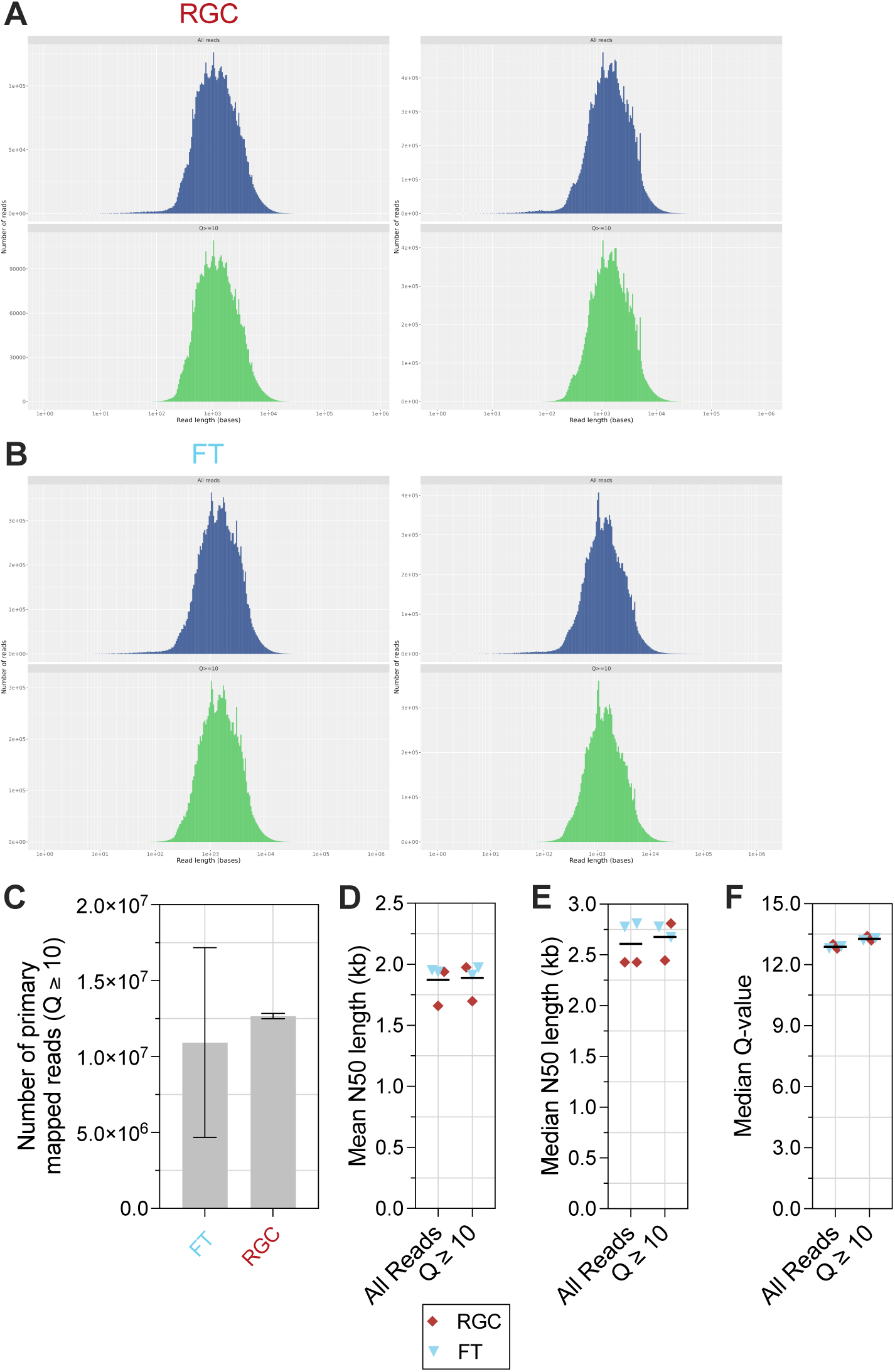
ONT sequencing of 2D purified RGCs and MACS flow-through (FT). **A-B)** Histograms showing read length distributions of all reads (top) and reads with a Q-score ≥ 7 (bottom) for **A)** RGC and **B)** FT replicates. **C)** Number of primary mapped ONT reads with a Q score ≥ 10 for RGCs and FT. Error bars denote standard deviation. **D)** Median N50 length for all reads. **E)** Mean N50 length of all reads and reads with a Q-score ≥ 10. **F)** Median Q-score for all reads and reads with Q ≥ 10. For D-F, black line denotes grand mean across all RGC and FT samples.

**Figure S15.**
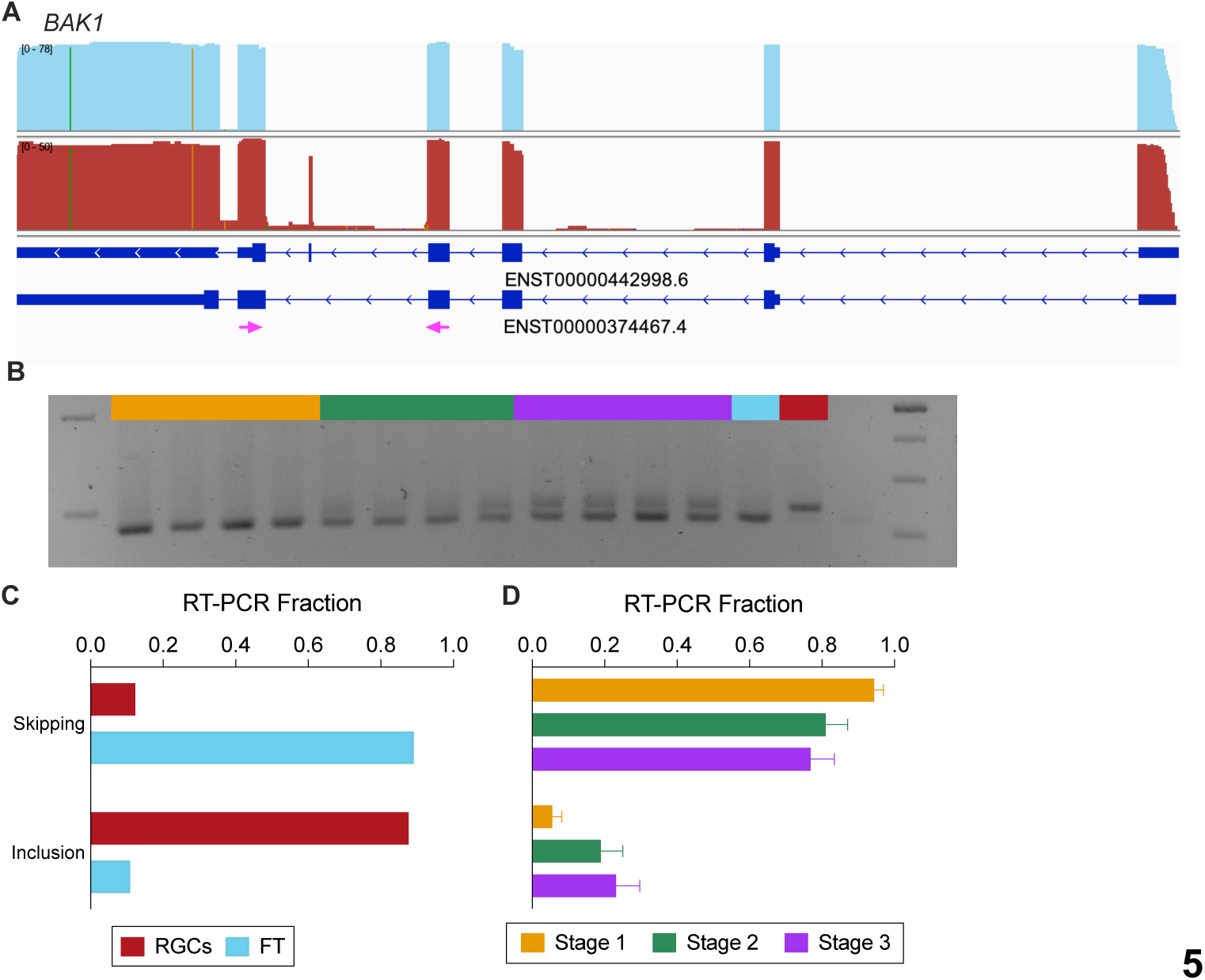
Validation of *BAK1* microexon splicing. **A)** IGV browser showing ONT reads mapping to the alternatively spliced microexon in RGCs but not FT. Pink arrows denote location of RT-PCR primers flanking the alternatively spliced microexon. **B)** Agarose gel image of *BAK1* RT-PCR amplicons. **C-D)** Quantification of *BAK1* microexon skipping and inclusion in **C)** RGCs and FT and **D)** retinal organoids. For D, error bars indicate SEM.

**Figure S16.**
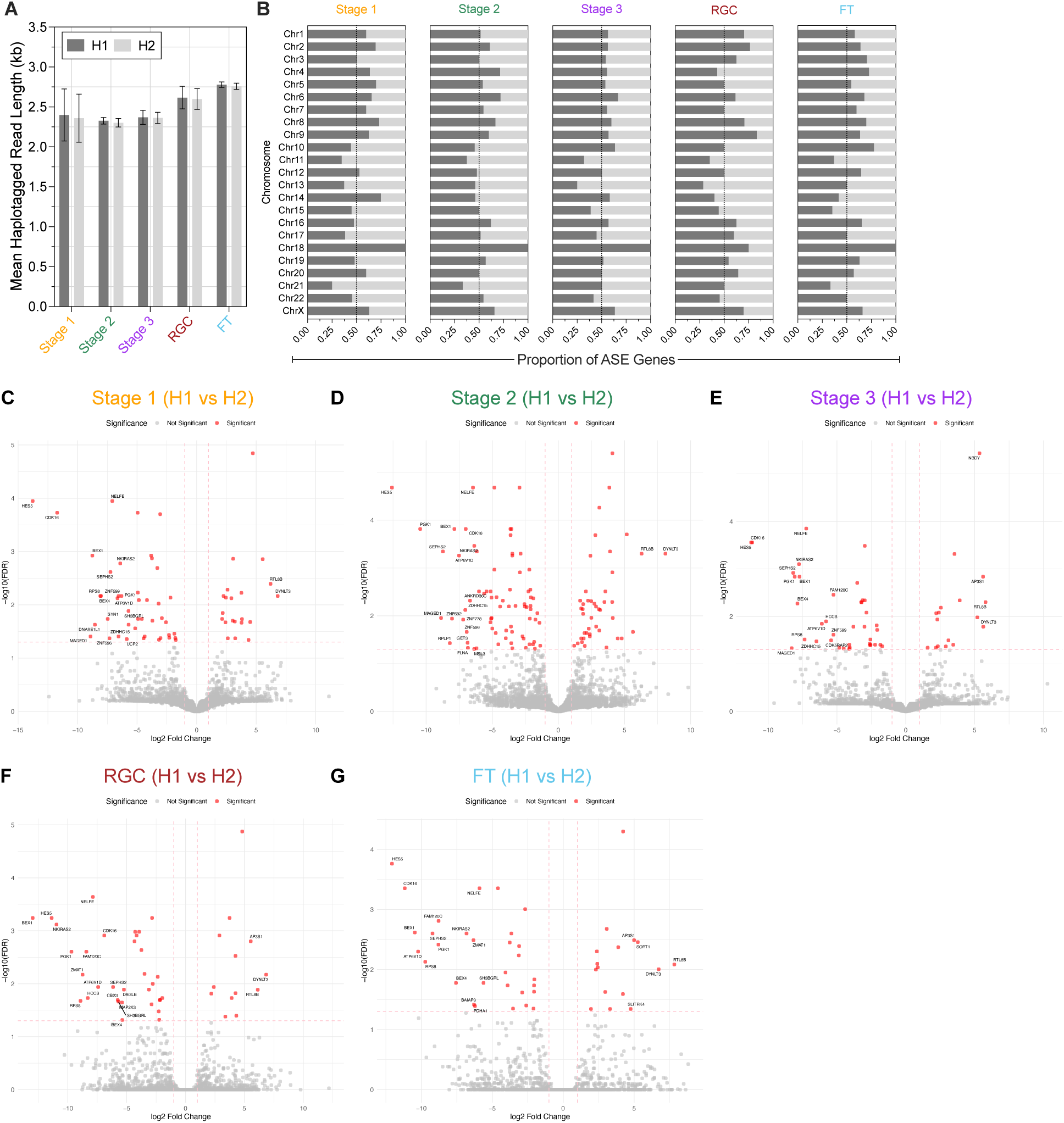
Allele-specific expression (ASE) analysis. **A)** Mean length of haplotagged ONT reads for each cell group. **B)** Proportion of ASE genes significant for either the H1 or H2 haplotype by chromosome, for each cell group. **C-G)** Volcano plots showing significant ASE (H1 vs H2, red dots) in **C)** Stage 1 organoids, **D)** Stage 2 organoids, **E)** Stage 3 organoids, **F)** 2D RGCs, and **G)** flow-through (FT).

**Figure S17.**
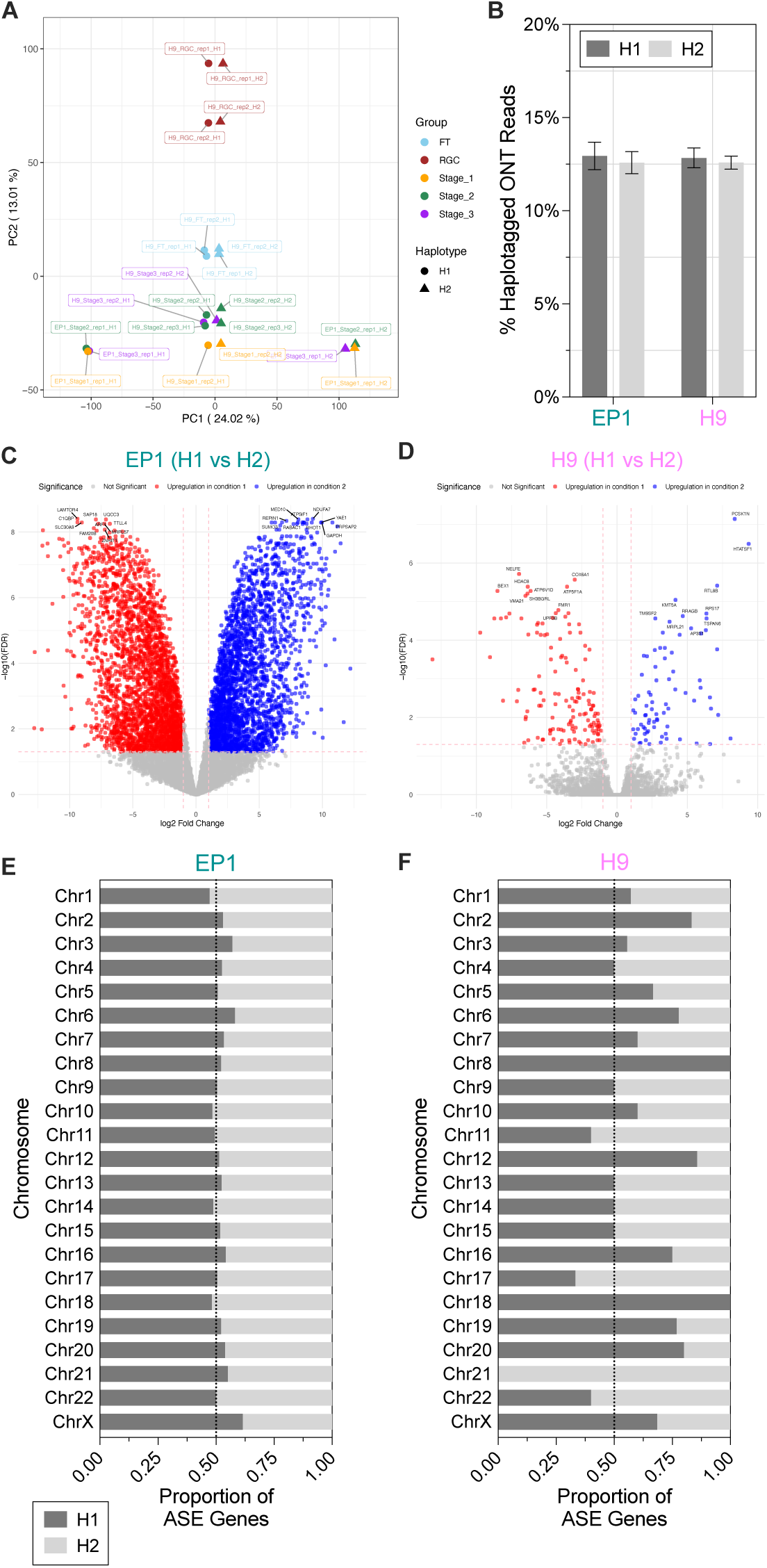
ASE in EP1- and H9-derived retinal organoids. **A)** PCA on phased gene counts. Color denotes group. Shape of dot indicates haplotype (H1 or H2). **B)** Percentage of haplotagged ONT reads from EP1 and H9 background organoids. Error bars denote standard deviation. **C-D)** Volcano plots showing allele-specific gene expression in **C)** EP1 and **D)** H9 background organoids. **E-F)** Proportion of ASE genes showing biases towards H1 or H2 in **E)** EP1 and **F)** H9 background organoid samples.

**Figure S18.**
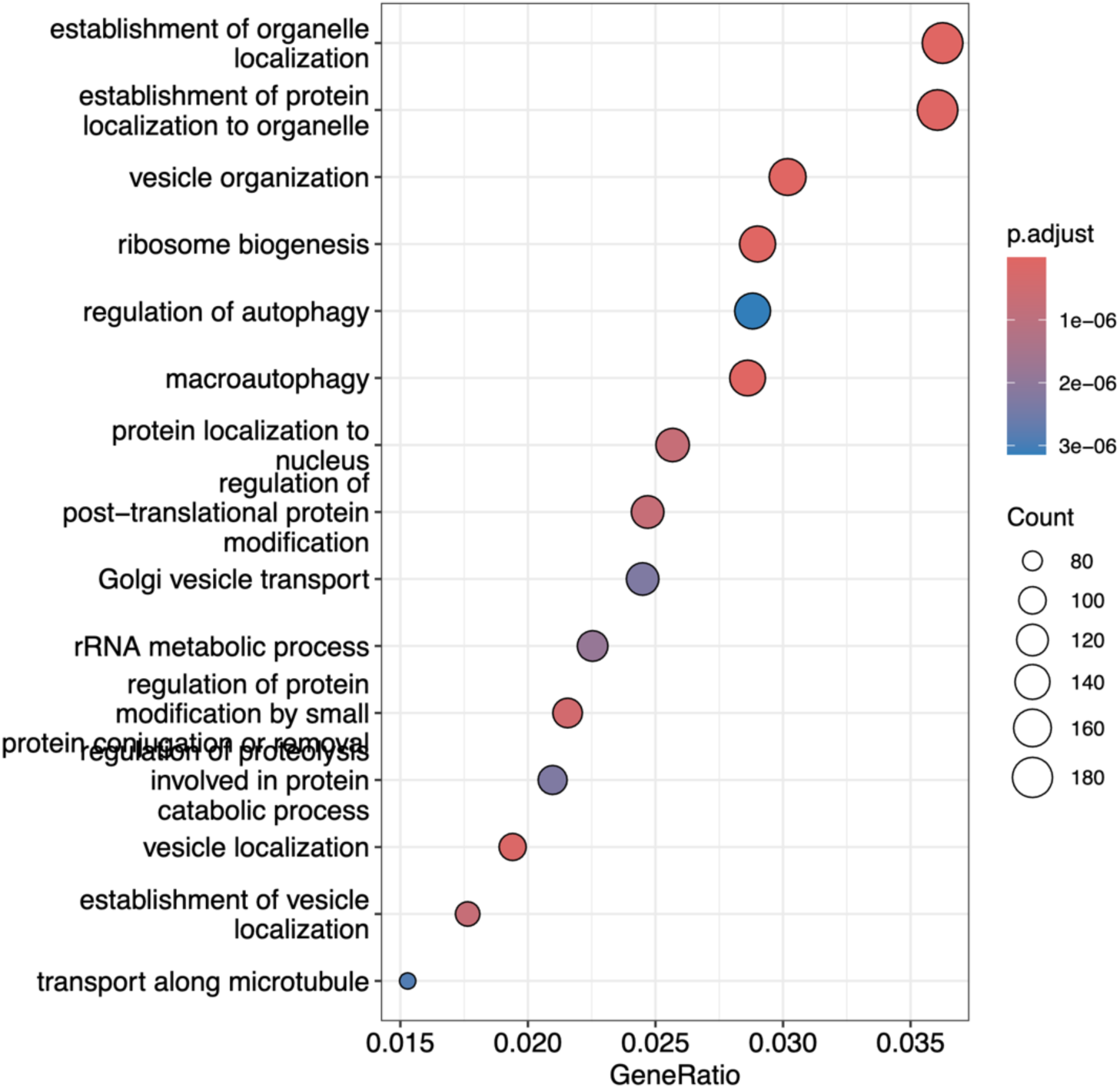
Gene Ontology (GO) analysis of gene with ASE in EP1-derived retinal organoid samples. Heatmap indicates adjusted p-value for enrichment, and dot size represents the number of significant genes found within each GO term.

**Figure S19.**
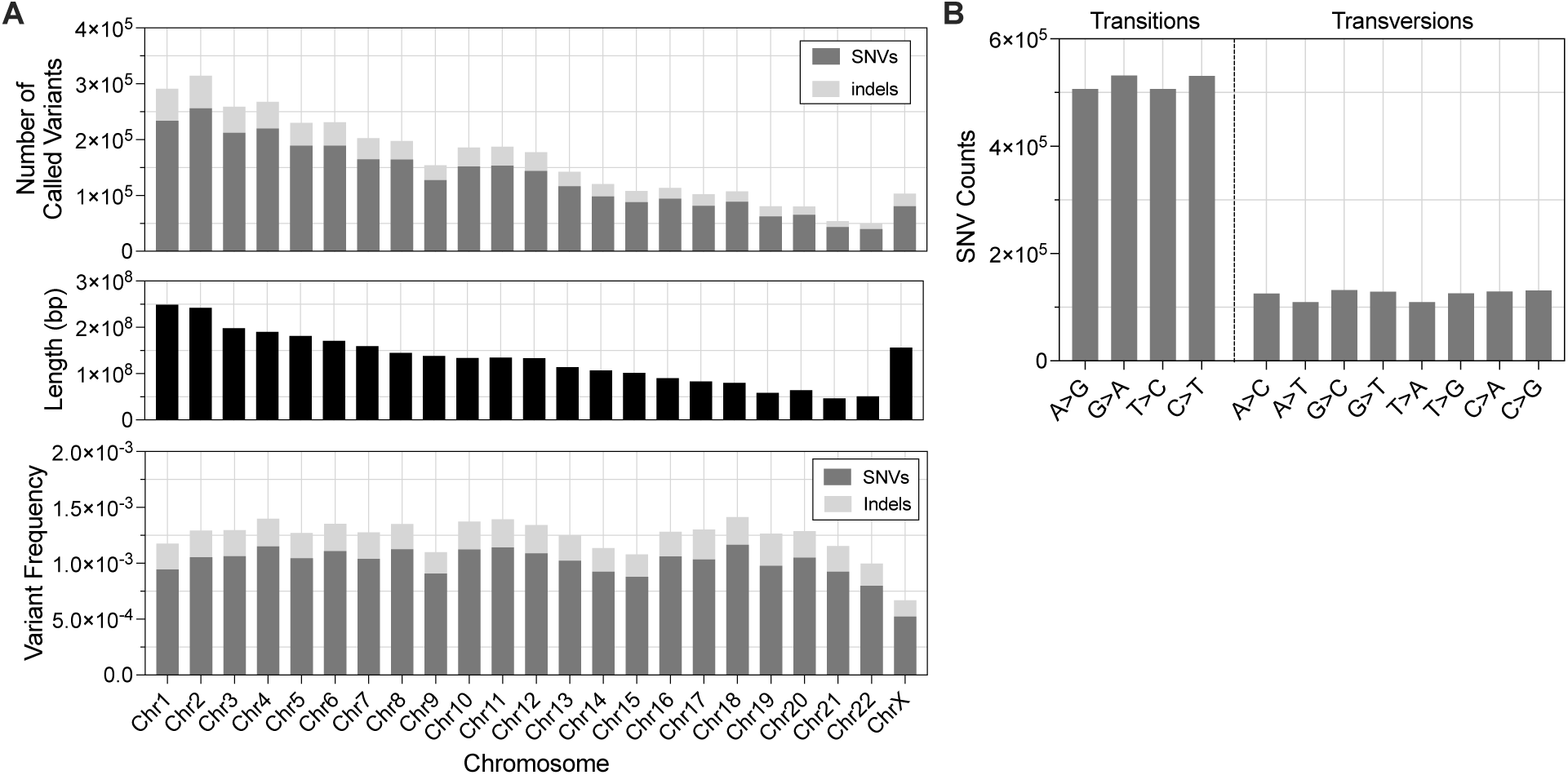
Variant Calling Format (VCF) file statistics. **A)** Number of called variants (SNVs and indels) in the VCF file by chromosome (top), with lengths of each chromosome (middle) and the variant frequency by chromosome (bottom). **B)** Number of each type of SNV in the VCF file.

**Figure S20.**
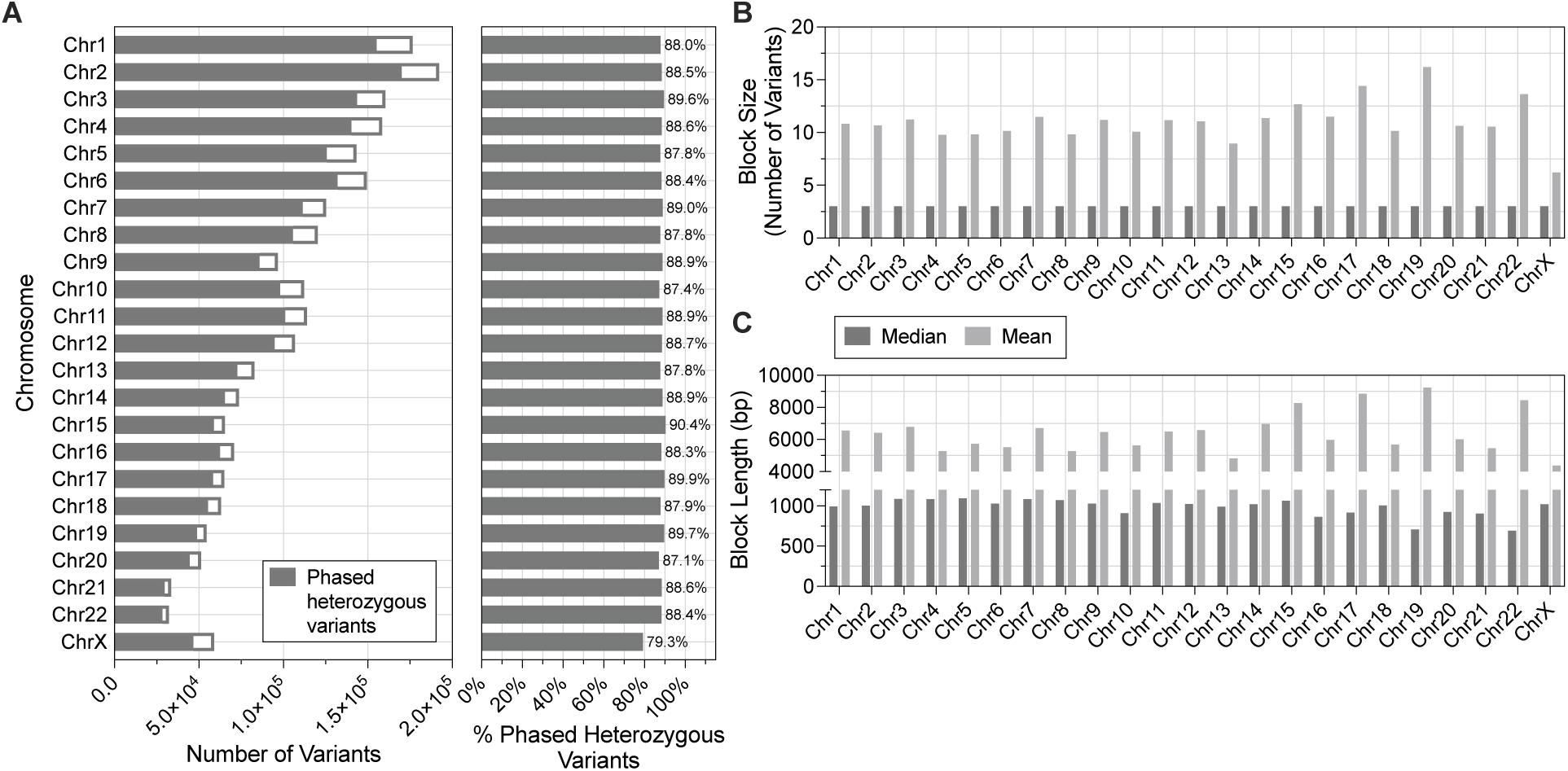
Phasing statistics. **A)** Number (left) and percentage (right) of phased heterozygous variants by chromosome. **B-C)** Average block (phase set) **B)** size and **C)** length by chromosome.

**Table S1.**
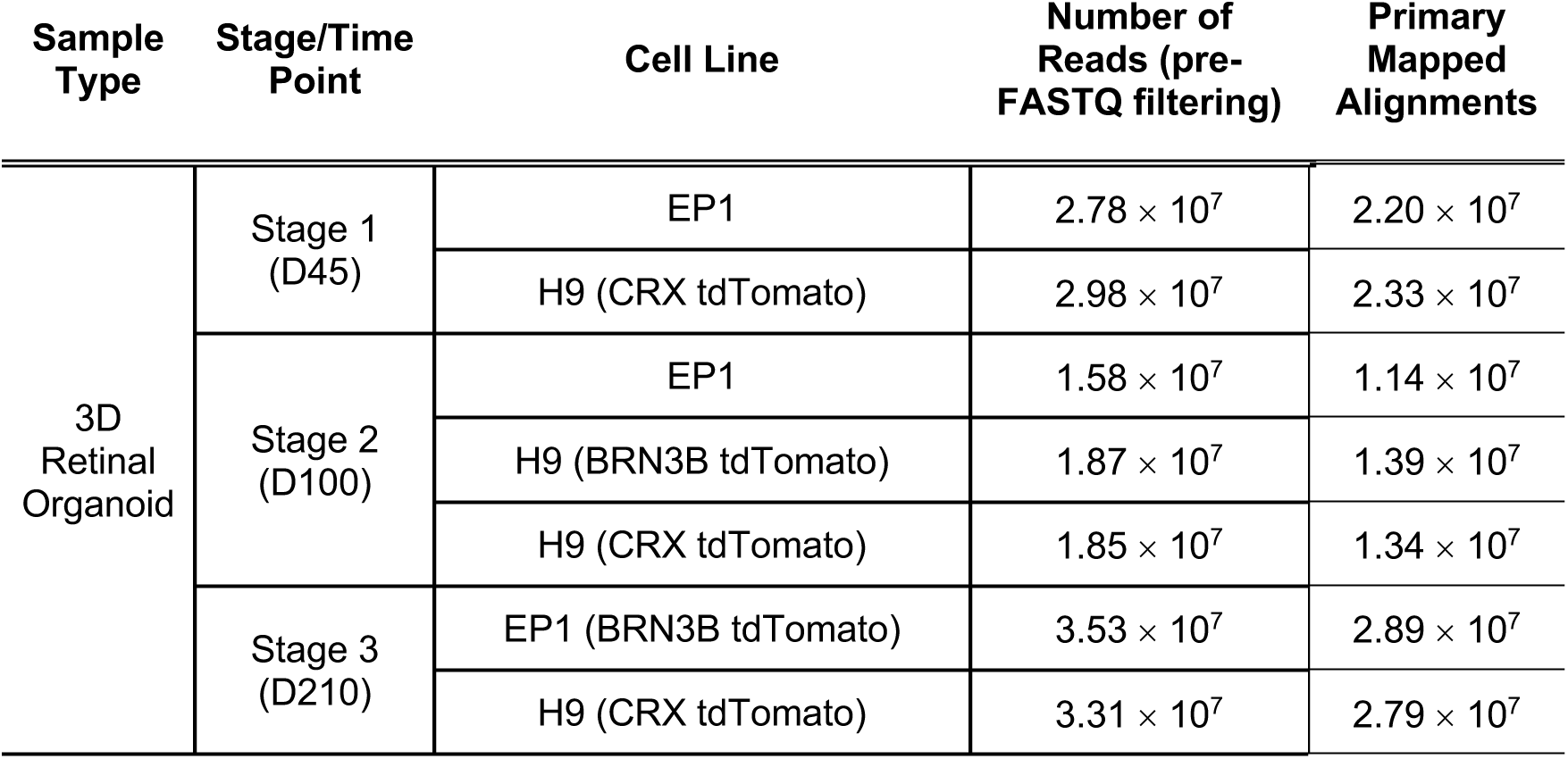
Summary of the direct ONT sequencing organoid samples in this study.

| Sample Type | Stage/Time Point | Cell Line | Number of Reads (pre-FASTQ filtering) | Primary Mapped Alignments |
| --- | --- | --- | --- | --- |
| 3D Retinal Organoid | Stage 1 (D45) | EP1 | $2.78 \times 10^7$ | $2.20 \times 10^7$ |
| | | H9 (CRX tdTomato) | $2.98 \times 10^7$ | $2.33 \times 10^7$ |
| | Stage 2 (D100) | EP1 | $1.58 \times 10^7$ | $1.14 \times 10^7$ |
| | | H9 (BRN3B tdTomato) | $1.87 \times 10^7$ | $1.39 \times 10^7$ |
| | | H9 (CRX tdTomato) | $1.85 \times 10^7$ | $1.34 \times 10^7$ |
| | Stage 3 (D210) | EP1 (BRN3B tdTomato) | $3.53 \times 10^7$ | $2.89 \times 10^7$ |
| | | H9 (CRX tdTomato) | $3.31 \times 10^7$ | $2.79 \times 10^7$ |

**Table S2.** ONT read statistics for retinal organoid samples (mean ± standard deviation).

|  |  | **Mean N50 Length (bp)** | **Median N50 Length (bp)** | **Median Q-value** |
| --- | --- | --- | --- | --- |
| Stage 1 | All Reads | 1706.9 ± 293.6 | 2326 ± 523.3 | 13.5 ± 0.1 |
| | Q ≥ 10 | 1686.1 ± 282.5 | 2254.5 ± 495.7 | 14.1 ± 0.1 |
| Stage 2 | All Reads | 1623.7 ± 103.5 | 2304.3 ± 204.9 | 11.9 ± 0.2 |
| | Q ≥ 10 | 1628 ± 94.6 | 2266 ± 197.6 | 12.7 ± 0.1 |
| Stage 3 | All Reads | 1724.8 ± 80.6 | 2329 ± 183.8 | 12.9 ± 0.1 |
| | Q ≥ 10 | 1743.2 ± 77.9 | 2337.5 ± 180.3 | 13.3 ± 0.1 |
| Combined | All Reads | 1676.4 ± 146.6 | 2317.6 ± 255.8 | 12.6 ± 0.7 |
| | Q ≥ 10 | 1677.5 ± 130.9 | 2283.1 ± 228.3 | 13.2 ± 0.6 |

**Table S3.** Differentially expressed genes (DEGs) in organoids across long-read (ONT) and short-read (Illumina) platforms (|log2FC| ≥ 1 & FDR < 0.05).

| Long Read Samples (ONT) | Short Read Samples (Illumina) | Number of Genes Tested | Total DEGs (ONT vs Illumina) | DEGs | Median GC of DEGs | Median transcript length of DEGs (bp) |
| --- | --- | --- | --- | --- | --- | --- |
|  |  |  |  | Up in ONT | Up in ONT | Up in ONT |
|  |  |  |  | Down in ONT | Down in ONT | Down in ONT |
| Stage 1 (D45) | D25 + D65 | 13,582 | 4,524 | 2,800 | 49.6% | 1,346 |
|  |  |  |  | 1,724 | 52.9% | 1,231 |
| Stage 2 (D100) | D65 + D100 | 13,783 | 5,063 | 3,209 | 49.6% | 1,449 |
|  |  |  |  | 1,854 | 53.8% | 1,235 |
| Stage 3 (D210) | D180 + D280 | 14,579 | 6,318 | 3,847 | 48.9% | 1,529 |
|  |  |  |  | 2,471 | 53.7% | 1,181 |

**Table S4.** Differentially expressed genes (DEGs) between retinal organoid stages (|log2FC| ≥ 1 & FDR < 0.05).

| Comparison | Total DEGs | Up DEGs |
| --- | --- | --- |
|  |  | Down DEGs |
| Stage 1 vs 2 | 1,220 | 730 |
|  |  | 490 |
| Stage 1 vs 3 | 4,144 | 2,573 |
|  |  | 1,571 |
| Stage 2 vs 3 | 1,633 | 994 |
|  |  | 639 |

**Table S5.**
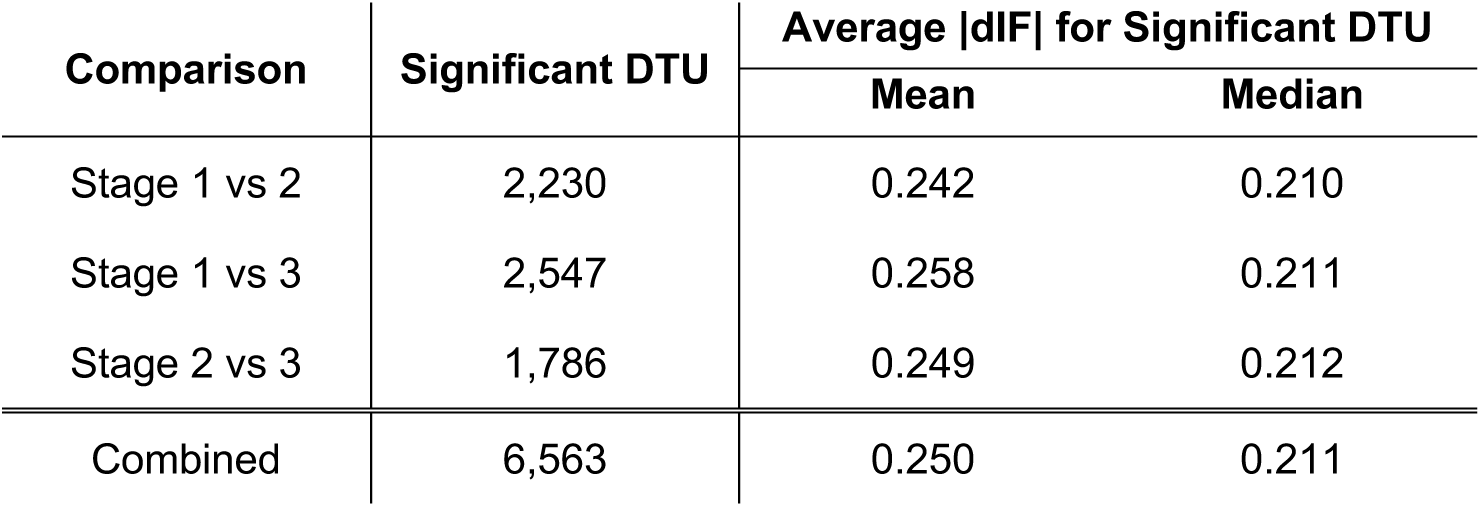
Mean and median dIF values for significant DTU events (|dIF| ≥ 0.1 & qval < 0.05) in retinal organoids.

**Table S6.** Chi-square testing for retinal disease (RD) genes (one-sided, at 95% confidence interval).

|  | RD Gene | Not RD Gene | Total | Chi-Square Test |  |
| --- | --- | --- | --- | --- | --- |
|  |  |  |  | Odds ratio | P-value |
| Multi-isoform Genes |  |  |  | 1.755 | 0.0001 |
| Multi-isoform | 162 | 6,961 | 7,123 |  |  |
| Single isoform | 56 | 4,224 | 4,280 |  |  |
| Total | 218 | 11,185 | 11,403 |  |  |
| Genes with Significant DTU |  |  |  | 1.587 | 0.0005 |
| Significant DTU | 82 | 3,079 | 3,161 |  |  |
| Not Significant DTU | 136 | 8,106 | 8,242 |  |  |
| Total | 218 | 11,185 | 11,403 |  |  |

**Table S7.**
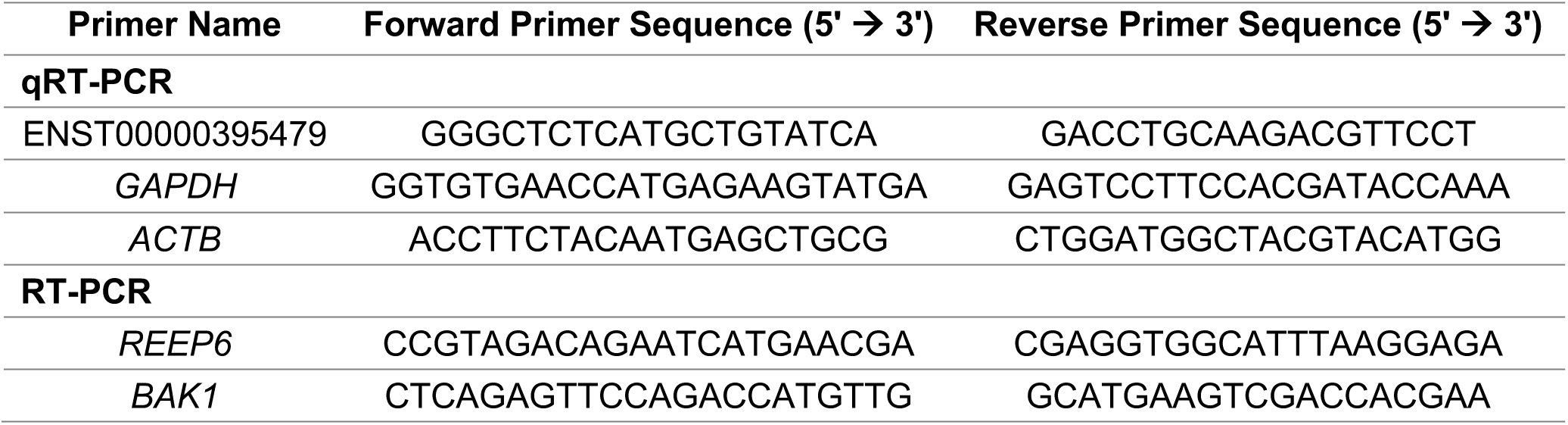
qRT-PCR/RT-PCR primers used in this study.

**Table S8.** ONT read statistics for 2D RGCs and flow-through (FT) samples (mean ± standard deviation).

|  |  | **Mean N50 Length (bp)** | **Median N50 Length (bp)** | **Median Q-value** |
| --- | --- | --- | --- | --- |
| RGC | All Reads | 1799.7 ± 196.9 | 2622 ± 275.8 | 12.9 ± 0.1 |
| | Q ≥ 10 | 1836.7 ± 196.4 | 2627.5 ± 258.1 | 13.3 ± 0.1 |
| FT | All Reads | 1945.3 ± 9.0 | 2791 ± 22.6 | 12.91 ± 0.1 |
| | Q ≥ 10 | 1941.4 ± 43.3 | 2725 ± 73.5 | 13.3 ± 0.1 |
| Combined | All Reads | 1872.5 ± 9.0 | 2706.5 ± 22.6 | 12.9 ± 0.1 |
| | Q ≥ 10 | 1889 ± 130.9 | 2676.3 ± 164.9 | 13.3 ± 0.1 |

**Table S9.** Shared ASE genes between all five retinal cell groups.

| Gene ID | Gene Name | Chromosome |
| --- | --- | --- |
| ENSG00000197921 | *HES5* | 1 |
| ENSG00000142937 | *RPS8* | 1 |
| ENSG00000003756 | *RBM5* | 3 |
| ENSG00000177879 | *AP3S1* | 5 |
| ENSG00000204356 | *NELFE* | 6 |
| ENSG00000124614 | *RPS10* | 6 |
| ENSG00000151893 | *CACUL1* | 10 |
| ENSG00000197345 | *MRPL21* | 11 |
| ENSG00000183955 | *KMT5A* | 12 |
| ENSG00000111775 | *COX6A1* | 12 |
| ENSG00000100554 | *ATP6V1D* | 14 |
| ENSG00000104129 | *DNAJC17* | 15 |
| ENSG00000179918 | *SEPHS2* | 16 |
| ENSG00000168256 | *NKIRAS2* | 17 |
| ENSG00000108953 | *YWHAE* | 17 |
| ENSG00000087111 | *PIGS* | 17 |
| ENSG00000152234 | *ATP5F1A* | 18 |
| ENSG00000181588 | *MEX3D* | 19 |
| ENSG00000102225 | *CDK16* | X |
| ENSG00000133169 | *BEX1* | X |
| ENSG00000102144 | *PGK1* | X |
| ENSG00000184083 | *FAM120C* | X |
| ENSG00000212747 | *RTL8B* | X |
| ENSG00000102409 | *BEX4* | X |
| ENSG00000165169 | *DYNLT3* | X |
| ENSG00000204272 | *NBDY* | X |
| ENSG00000189369 | *GSPT2* | X |

**Table S10.**
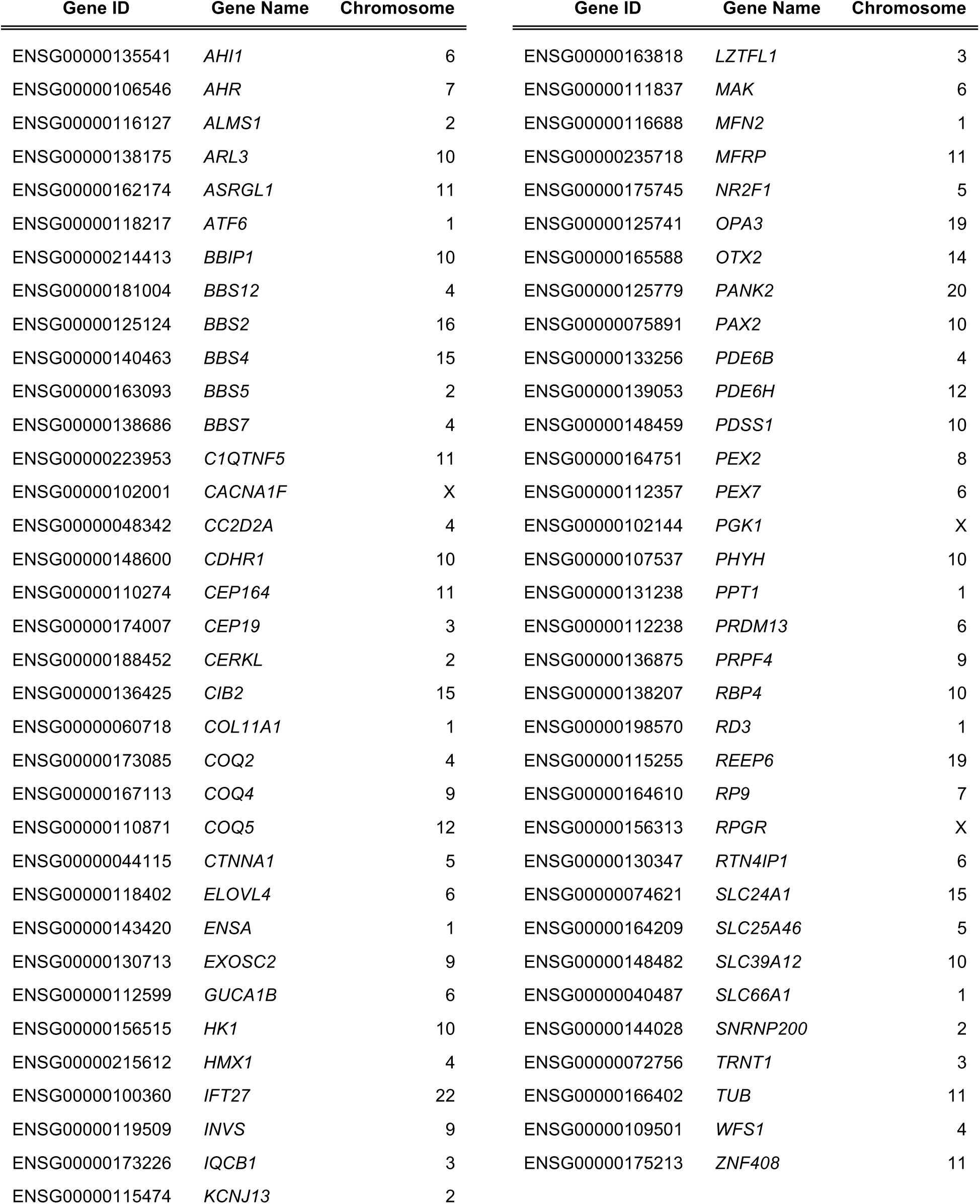
Retinal disease-associated genes with ASE in EP1-derived retinal organoids.

| Gene ID | Gene Name | Chromosome | Gene ID | Gene Name | Chromosome |
| --- | --- | --- | --- | --- | --- |
| ENSG00000135541 | *AHI1* | 6 | ENSG00000163818 | *LZTFL1* | 3 |
| ENSG00000106546 | *AHR* | 7 | ENSG00000111837 | *MAK* | 6 |
| ENSG00000116127 | *ALMS1* | 2 | ENSG00000116688 | *MFN2* | 1 |
| ENSG00000138175 | *ARL3* | 10 | ENSG00000235718 | *MFRP* | 11 |
| ENSG00000162174 | *ASRGL1* | 11 | ENSG00000175745 | *NR2F1* | 5 |
| ENSG00000118217 | *ATF6* | 1 | ENSG00000125741 | *OPA3* | 19 |
| ENSG00000214413 | *BBIP1* | 10 | ENSG00000165588 | *OTX2* | 14 |
| ENSG00000181004 | *BBS12* | 4 | ENSG00000125779 | *PANK2* | 20 |
| ENSG00000125124 | *BBS2* | 16 | ENSG00000075891 | *PAX2* | 10 |
| ENSG00000140463 | *BBS4* | 15 | ENSG00000133256 | *PDE6B* | 4 |
| ENSG00000163093 | *BBS5* | 2 | ENSG00000139053 | *PDE6H* | 12 |
| ENSG00000138686 | *BBS7* | 4 | ENSG00000148459 | *PDSS1* | 10 |
| ENSG00000223953 | *C1QTNF5* | 11 | ENSG00000164751 | *PEX2* | 8 |
| ENSG00000102001 | *CACNA1F* | X | ENSG00000112357 | *PEX7* | 6 |
| ENSG00000048342 | *CC2D2A* | 4 | ENSG00000102144 | *PGK1* | X |
| ENSG00000148600 | *CDHR1* | 10 | ENSG00000107537 | *PHYH* | 10 |
| ENSG00000110274 | *CEP164* | 11 | ENSG00000131238 | *PPT1* | 1 |
| ENSG00000174007 | *CEP19* | 3 | ENSG00000112238 | *PRDM13* | 6 |
| ENSG00000188452 | *CERKL* | 2 | ENSG00000136875 | *PRPF4* | 9 |
| ENSG00000136425 | *CIB2* | 15 | ENSG00000138207 | *RBP4* | 10 |
| ENSG00000060718 | *COL11A1* | 1 | ENSG00000198570 | *RD3* | 1 |
| ENSG00000173085 | *COQ2* | 4 | ENSG00000115255 | *REEP6* | 19 |
| ENSG00000167113 | *COQ4* | 9 | ENSG00000164610 | *RP9* | 7 |
| ENSG00000110871 | *COQ5* | 12 | ENSG00000156313 | *RPGR* | X |
| ENSG00000044115 | *CTNNA1* | 5 | ENSG00000130347 | *RTN4IP1* | 6 |
| ENSG00000118402 | *ELOVL4* | 6 | ENSG00000074621 | *SLC24A1* | 15 |
| ENSG00000143420 | *ENSA* | 1 | ENSG00000164209 | *SLC25A46* | 5 |
| ENSG00000130713 | *EXOSC2* | 9 | ENSG00000148482 | *SLC39A12* | 10 |
| ENSG00000112599 | *GUCA1B* | 6 | ENSG00000040487 | *SLC66A1* | 1 |
| ENSG00000156515 | *HK1* | 10 | ENSG00000144028 | *SNRNP200* | 2 |
| ENSG00000215612 | *HMX1* | 4 | ENSG00000072756 | *TRNT1* | 3 |
| ENSG00000100360 | *IFT27* | 22 | ENSG00000166402 | *TUB* | 11 |
| ENSG00000119509 | *INVS* | 9 | ENSG00000109501 | *WFS1* | 4 |
| ENSG00000173226 | *IQCB1* | 3 | ENSG00000175213 | *ZNF408* | 11 |
| ENSG00000115474 | *KCNJ13* | 2 |  |  |  |

**Table S11.** ASE analysis in EP1-derived retinal organoids of mapped and named loci associated with Age-related macular degeneration (AMD).

| **Gene Name** | **Gene ID** | **Chromosome** | **Log2FC** | **PValue** | **Adj PValue** |
| --- | --- | --- | --- | --- | --- |
| ***HTRA1*** | **ENSG00000166033** | **10** | **-3.841404683** | **3.07E-05** | **1.89E-04** |
| *CFH* | ENSG00000000971 | 1 | -0.02038222 | 9.91E-01 | 9.97E-01 |
| *CFB* | ENSG00000243649 | 6 | -5.337876486 | 2.35E-02 | 5.49E-02 |
| *C3* | ENSG00000125730 | 19 | -1.928638151 | 1.98E-01 | 3.10E-01 |

